# HDX-MS reveals concealed conformations of ISWI during different stages of nucleosome sliding

**DOI:** 10.1101/2023.07.30.549839

**Authors:** Younus A Bhat, Javaid Y Bhat, Shajrul Amin, Jayant B Udgaonkar, Ajazul H Wani

**Author notes:** Any correspondence can be addressed to Ajaz ul H Wani http://biotechnology.uok.edu.in/Main/ViewPage.aspx?Page=d8376142-bef2-4eec-890c-64dade6c4130.

## Abstract

Nucleosome spacing across the genome is regulated by the adenosine 5’-triphosphate (ATP)- dependent nucleosome sliding activity of Imitation Switch, ISWI. ISWI is believed to be auto-inhibited in the resting-state by the binding of its N-and C-terminal regulatory regions to its central ATPase-domain, attaining a “closed” conformation. To slide nucleosomes ISWI must i) transition to the state competent for nucleosome binding, ii) bind to nucleosome and iii) carry the ATP-dependent nucleosome sliding. The conformations attained by full-length ISWI (FL-ISWI) during the entire sliding process have remained inaccessible by the methods used so far. Using Hydrogen/Deuterium-exchange coupled to Mass-Spectrometry (HDX-MS), we monitored conformational dynamics of the *Drosophila* FL-ISWI at all the stages of sliding process. HDX-MS data show that in the resting state, ISWI samples an ensemble of conformations showing varying levels of deuterium uptake in many regions including N-and C-terminal regulatory regions, suggesting ISWI intrinsically samples relatively “open-states”. In addition to substantiating previous nucleosome binding studies, HDX-MS reveals that during actual sliding-step, regions of ATPase-domain which bind to the nucleosomal DNA undergo major conformational change. The C-terminal HSS domain switches from the solvent protected stable state to a more dynamic state, implying several interactions established by ISWI with the nucleosome upon binding are relieved during sliding. In sum, this study provides mechanistic insights into how ISWI can switch from an auto-inhibited “closed-state” to an “open-state” competent for nucleosome binding, and reveals the conformation attained by ISWI during the actual nucleosome sliding step. We propose that, like ISWI, intrinsic dynamics may be involved in functioning of other Rec-like ATPase-domain containing protein families.

## Introduction

Multi-scale folding of chromatin from nucleosomes to chromosomes ensures packaging of chromatin within the nucleus (1). At the primary level, 147 bps of DNA are wrapped around an octamer of histone proteins forming a nucleosome (2). Nucleosomes are formed all along the genome, but their assembly, disassembly, occupancy and accessibility, as well as inter-nucleosome spacing, is highly regulated by enzymatic and non-enzymatic processes (3-6). ATP-dependent chromatin remodelers utilize the energy from ATP hydrolysis to remodel nucleosomes and regulate their assembly, disassembly and spacing (3,7,8,9). Perturbations in the functioning of chromatin remodelers have been linked with different diseases (4,10), necessitating to understand the regulation and mechanism of action of these remodelers.

Chromatin remodelers are classified into four main groups SWItch/Sucrose Non-Fermentable (SWI/SNF), Chromodomain-Helicase-DNA binding (CHD), Inositol requiring 80 (INO80) and Imitation switch (ISWI). All of them share a conserved ATPase domain consisting of two Rec-A like domains (core-1 and core-2) but differ in flanking regions (4). The ISWI family of remodelers is a structurally conserved family of ATPases with three main domains, the N-terminal region (NTR) (containing AutoN, ppHSA (post-post-helicase-SANT-associated) domains), the ATPase domain (consisting of core-1 and core-2) and the C-terminal NegC region and HSS (HAND-SANT-SLIDE) domain (11,12) (Figure 1 A, B). AutoN, ppHSA and NegC have been shown to negatively regulate the activity of the ATPase domain, while HSS domain is involved in binding to the extra-nucleosomal DNA (11,12,13,14).

**Figure 1.**
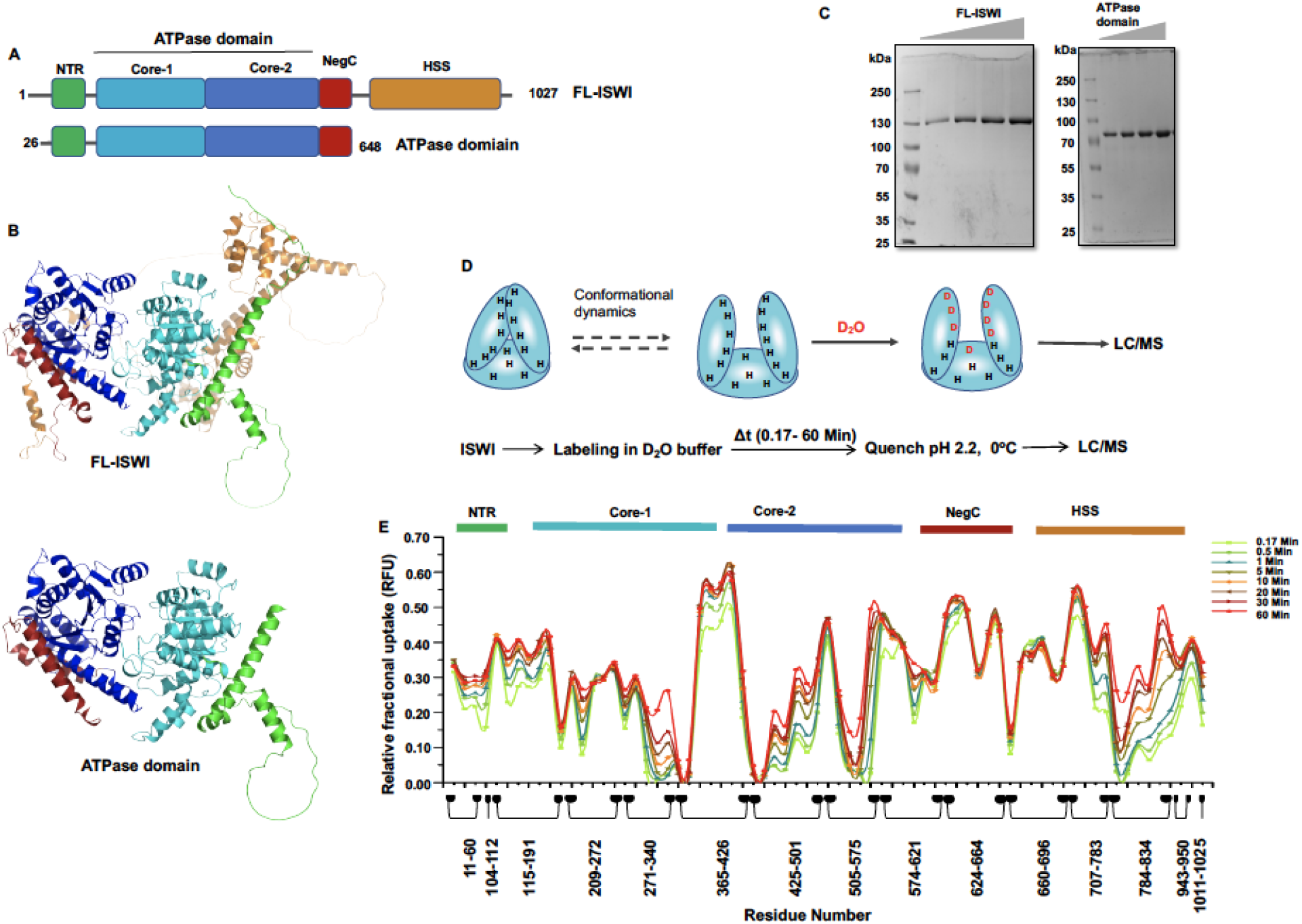
Intrinsic conformational dynamics of ISWI: (A) Domain organization of full-length ISWI (FL-ISWI) and its ATPase domain. (B) AlphaFold (30,31) structure of *Drosophila* ISWI(ID:AF-Q24368-F1) showing the N-terminal region (green), Core-1 of ATPase domain (cyan), core-2 of ATPase domain (dark blue), NegC (dark-red) and C-terminal (brown) regions. (C) SDS-PAGE showing purified ISWI-FL and ATPase domain. (D) Schematic showing conformational dynamics of a protein can result in transient exposure of some exchangeable hydrogens to the solvent which can get exchanged with solvent deuteriums if D_2_O is in excess and number of deuteriums incorporated can be detected by LC-MS. (Below) Flow chart showing the main steps of the HDX-MS protocol followed. (E) Relative fractional uptake of deuterium (RFU) of FL-ISWI at varying labelling times. The multiple colored plots show different labelling times (0.17 min to 60 min). The numbers on x-axis reflect the first and last residues of the different regions across the ISWI sequence.

*In vitro* reconstituted nucleosome remodeling systems have proven very useful in understanding the mechanism of nucleosome sliding (11-19). The ATPase domain of ISWI is an autonomous remodeling unit but its activity is tightly regulated by its N-and C-terminal domains (Fig. 1A, B) (11, 13, 17, 22). In the absence of nucleosomes, the NTR docks on the ATPase domain and negatively regulates its ATPase activity. But it gets dislodged when histone H4-tail binds to the ATPase domain near the same region where NTR binds. The NegC domain also binds to the ATPase domain and inhibits the ATPase enzyme activity, with mechanistic details still obscure. In brief, ISWI seems to attain an “auto-inhibited” state in the absence of nucleosomes, but in the presence of nucleosomes, ISWI has to overcome this auto-inhibition and interact with nucleosomal epitopes, the H4-tail and extra-nucleosomal DNA (18). This mechanism is conserved from yeast to humans (11, 17,18, 22). But how ISWI switches from an auto-inhibited state to a nucleosome-binding competent state is not clear.

Conformational changes in the ATPase domain of ISWI upon binding to the nucleosome and/or ATP analogues have been analyzed by cryo-EM and X-ray crystallography (17, 20, 21). Change in orientation of ATPase core-1 and core-2 domains were observed upon nucleosome binding (17,21). However, C-terminal regions were not detected in the nucleosome bound state (17,21). These altered conformations have been proposed to represent the activated/primed state of ISWI. Despite the availability of X-ray or cryo-EM structures of the ISWI ATPase bound to either nucleosome and/or ATP analogues, the conformations attained by ISWI during the actual nucleosome sliding step could not be revealed. However, to obtain a proper understanding of the mechanism of chromatin remodeling, it is necessary to simultaneously monitor conformational changes in all the domains of ISWI during the actual nucleosome sliding process. Studying the conformational dynamics of ISWI during nucleosome sliding has proven to be challenging and has not been achieved.

To monitor the conformational dynamics of full-length ISWI in its resting-state (native/apo-state) and during actual nucleosome sliding step we employed a high-sensitivity method, Hydrogen/Deuterium Exchange coupled to Mass Spectrometry (HDX-MS). Exchangeable hydrogens (N-, O-and S-linked) of a protein get replaced by solvent deuterons when a protein undergoes local or global unfolding event, or a conformational change and the sites of exchange become accessible to the solvent (23-26). Even under native conditions proteins are dynamic structures undergoing intrinsic thermal fluctuations of varying magnitude (27, 28). These fluctuations enable proteins to sample different conformational sub-states under native conditions (28). Therefore, proteins under native conditions are not in a unique conformation but exist as an ensemble of conformations (27-30). The fraction of molecules in different conformational sub-states is governed by Boltzmann distribution (i.e. the free-energy difference between any sub-state and the most stable state) (31). Most stable conformation is highest populated while least stable conformation is lowest populated but molecules exchange/shuttle between different conformational sub-states. HDX-MS, can detect these different conformational sub-states even if sampled transiently by a protein (Fig. 1D) (21-26). The exchange of protein molecules between different conformations differing in solvent exposure of even one or more exchangeable hydrogen makes them detectable by HDX-MS (Fig. 1D). We and others have shown by HX-MS and by HX-NMR that under native conditions proteins are in equilibrium with different partially folded or even fully unfolded conformations *(24,32-40)*. Furthermore, this method can be used to monitor conformational transitions in time scale of milliseconds to hours and possibly with protein of any size (25, 41-43).

Here, we report HDX-MS data that clearly show ISWI undergoes intrinsic conformational fluctuations in the “resting-state” sampling an ensemble of conformations with auto-inhibitory regions (N-and C-terminal regions) interacting with ATPase-core as “closed” state or displaced from the ATPase core as “open” state, thereby, providing a transient opportunity for ISWI-nucleosome interactions. During nucleosome sliding, ISWI undergoes a global conformational change in comparison to nucleosome bound-state encompassing all the major domains. These data provide molecular insights into how ISWI switches from the auto-inhibited state to the nucleosome-binding competent state as well as the conformational changes it undergoes in order to transition from the nucleosome-bound to nucleosome-sliding state. Our analysis also suggests a possible connection between intrinsic and sliding conformations of ISWI.

## Results

### Resting-state conformational dynamics of FL-ISWI

ISWI has been proposed to exist in an “auto-inhibited” resting-state with N-and C-terminal regions interacting with its ATPase domain (11, 13, 17, 22), now on referred as “closed” state. In the presence of nucleosome, the N-and C-terminal regions are dislodged from the ATPase core for interaction with the nucleosome. Given the coincident binding sites of its N-and C-terminal regions (auto-inhibitory regions) and nucleosomal epitopes on the ATPase domain, this mechanism demands that ISWI has to overcome the auto-inhibition in order to bind to the nucleosome. We hypothesized that ISWI samples an “open” state in which the N-and C-terminal regions get transiently displaced from ATPase domain thereby providing a time window for ISWI-nucleosome interactions. To test our hypothesis, we monitored the intrinsic conformational dynamics of full length ISWI (FL-ISWI) (Fig. 1A, B, C) by HDX-MS in its resting state (ISWI alone). HDX-MS can detect these conformational sub-states even if sampled transiently by ISWI, because “open” conformation will exchange more number of deuteriums than “closed” conformation, and hence, will have higher mass (Fig. 1D) (34-39). To carry HDX-MS, FL-ISWI was diluted with D_2_O labeling buffer and incubated for increasing time durations (10s to 1h). Exchange was quenched, and the protein was pepsin digested. The extent of time-dependent deuterium uptake in different regions was determined by LC-MS (Fig. 1D). We analyzed only peptic peptides which were detected across all experiments and labelling times (71 peptides) covering about 60% of the protein with a redundance of 1.55 across different states (see methods) (Fig. S18). We observed mass spectra gradually shifting to higher mass with increasing labeling time having unimodal peak distribution, suggesting that exchange occurs via local opening conformational fluctuations or EX2 mechanism (Fig. S1). Relative deuterium uptake was plotted across the ISWI sequence for different labeling times (Fig. 1E). Based on the level of deuterium uptake (Fig. 1E), different regions of ISWI appear to fall in three major categories: a) regions showing no significant deuterium uptake (Class-a), b) regions showing deuterium uptake at the first time point of labeling but no further increase with increasing labeling time (Class-b) and c) regions showing increase in deuterium uptake with increase in labeling time (Class-c) (Sup. Table-1). Exchangeable hydrogens get replaced by solvent deuterons when a protein undergoes local or global unfolding event and the sites of exchange become accessible to the solvent (see methods) (23,24,25,26). Therefore, Class-a regions do not seem to fluctuate sufficiently to attain an exchange competent state for H/D exchange to occur within the time scale of labeling and hence, remain protected. The Class-b regions undergo local unfolding fluctuations sufficient to attain an exchange competent state in which only some of the residues of each region get exposed to the solvent while the rest remain protected, resulting in no further deuterium uptake with increasing labeling time. The Class-c regions undergo more and more unfolding fluctuations within the labeling time scale, incorporating more and more deuterons and hence, follow a time-dependent deuterium uptake (Fig. 1E, S1, S2, Table-S1). The three classes of regions seem to have different protection factors and hence their deuterium uptake levels are proportional to the time of labeling.

**Figure 2.**
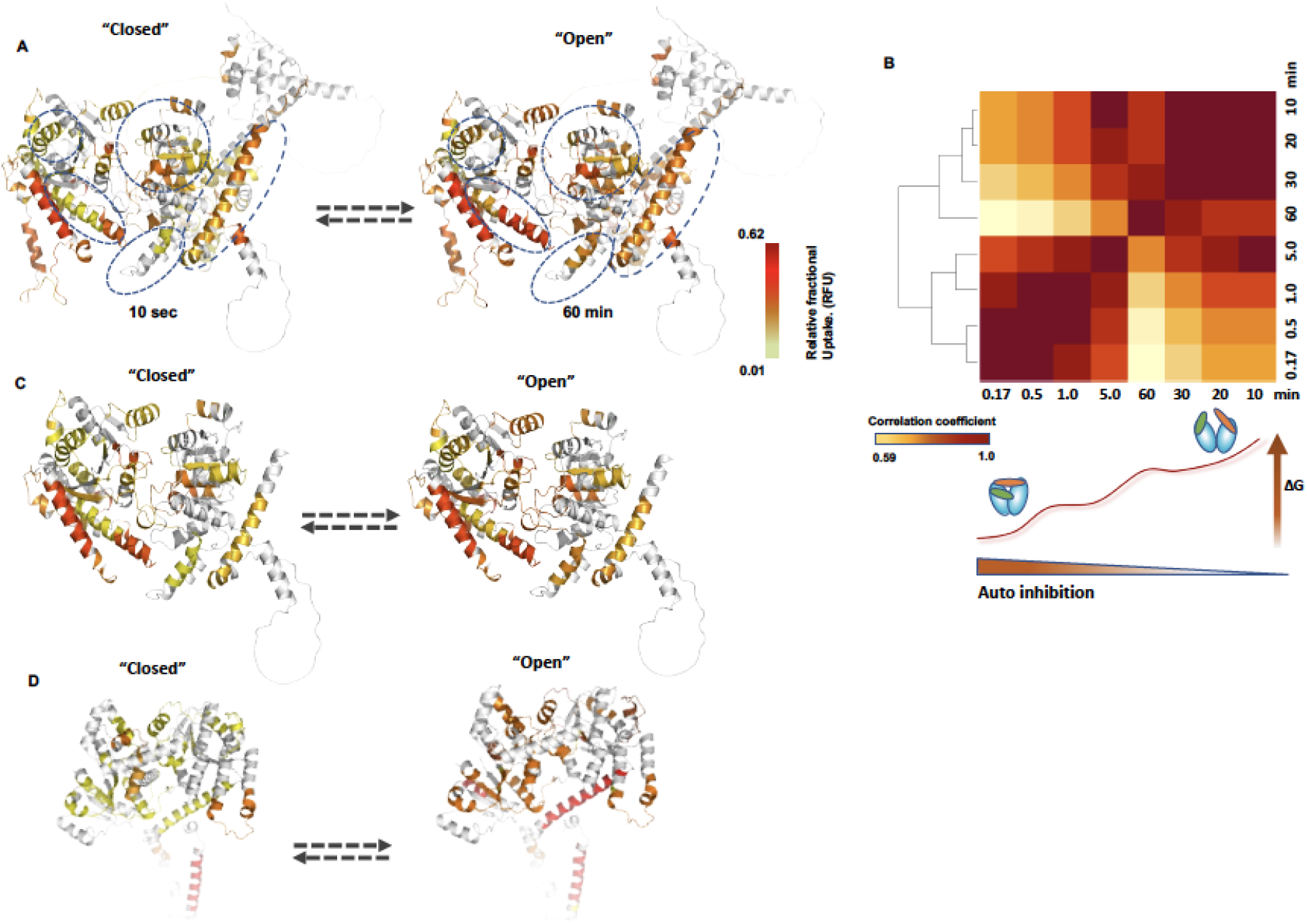
Intrinsic conformational ensemble of ISWI: Deuterium uptake levels mapped on FL-ISWI (ID:AF-Q24368-F1) (A) ATPase domain (C) and on MtISWI ATPase domain (PDB code 5JXR) (D) at 10s (closed) or 1 h (open) of labelling. Dashed ovals highlight some regions showing an increase in deuterium uptake with increasing labelling time (the class-c regions). Regions with no peptide coverage are colored gray. (B) Clustering of conformations sampled by ISWI. Schematic showing open and closed conformations of ISWI. Green: NTR, Blue: ATPase core, Orange: C-terminal region.

As no full-length structure of ISWI is available, the Alpha fold structure (ID:AF-Q24368-F1) (44, 45) was used to depict the structural changes in FL-ISWI (Fig. 1B). Mapping of the time-dependent deuteration of these regions on to the FL-ISWI Alpha fold structure shows that ISWI samples multiple conformations that differ in solvent accessibility at different regions and hence, in the deuterium uptake levels (Fig. 2A, S3). Conformations which take up less deuterons are relatively folded (closed) while as conformations which take up more deuterons are partially unfolded (open) and more accessible to solvent. However, these data cannot report on the rate of conformational dynamics or exact number of conformations sampled by ISWI or the order in which they are sampled as the exchange usually occurs by EX2 mechanism under native conditions (see methods). Conformations detected at different labeling times were correlated with each other by comparing the deuteration of corresponding regions (Fig. S4). Based on the correlation values, conformations of ISWI cluster into two main groups, depending on similarity in deuteration (Fig. 2B). Conformations with lower deuteration (at shorter labeling times) represent a relatively folded “closed” state, while conformations with a higher extent of deuteration represent an “open” state (Fig. 2B). Many regions from the NTR, core-1 and core-2 interface, NegC and the HSS domain show an increase in deuteration with time (Class-c) (Fig. 1E, Fig. 2A, Fig. S2, S3), implying that these regions transiently attain a solvent accessible open state. Most of the Neg-C and the adjacent region towards the HSS domain show more than 35% of deuteration at just 10s of labeling, implying it is solvent accessible or in a highly dynamic state (Fig. 1E, 2A). The interaction of the NTR and Neg-C with the ATPase core is believed to impose auto-inhibition of ISWI (11,13,17,22). But inherent conformational fluctuations of these regions between the lower and higher deuterated states (Fig. 1E, Fig. S2) suggests that ISWI samples both “closed” and “open” conformational sub-states in which these regions interact with ATPase domain and are protected or get dislodged from ATPase domain resulting in H/D exchange, respectively. Whether NegC masks the ATPase core or is protruded out is still not resolved (12,17). The observation of time-dependent increase in deuteration of NegC suggests that NegC is not stitched to ATPase domain but possibly fluctuates, and hence, could explain both of the previous observations where NegC was reported to exist in either the ATPase proximal or protruded conformations.

Mapping of the deuterium uptake on the crystal structure of *M. thermophila* ISWI (MtISWI, residues 81–723) (17) after 10 s and 1 h of labelling (to represent lower and higher deuterated states) shows that the NTR as well as the ATPase domain show differential deuterium uptake at these two labelling times (Fig. 2D). MtISWI has been shown to attain an “auto-inhibited” state in which the NTR is bound to the ATPase domain. Uptake of more deuterium at 1 h of labelling compared to at 10s suggests intrinsic fluctuations of ISWI between relatively “closed” and “open” conformational substates with more or less auto-inhibition, respectively (Fig. 2C). Overall, these results suggest that FL-ISWI intrinsically samples multiple interchangeable conformations with varying solvent exposure and possibly having varying degrees of auto-inhibition (Fig. 2A, B, S3).

**Figure 3.**
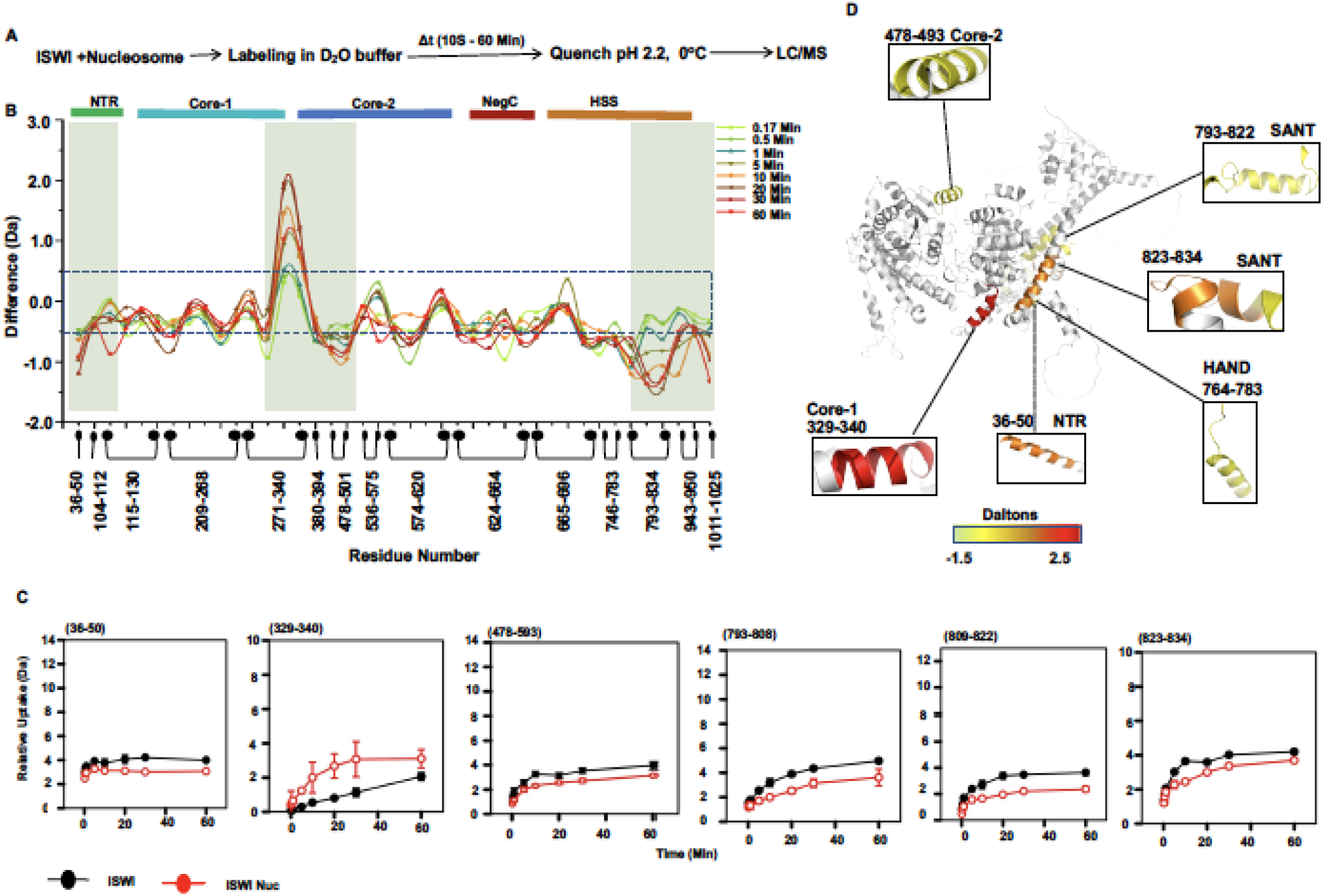
Conformation attained by ISWI upon nucleosome binding. (A)Flow chart showing main steps of the HDX-MS experiment for nucleosome binding. (B) Difference (subtraction) plot of ISWI-Nucleosome and ISWI at different labelling times. Differences in deuterium exchange of more than ±0.5 Da are considered significant (dashed box). Shaded regions show significant change. (C) Regions of ISWI showing significant differential deuterium uptake in the presence and absence of nucleosome. (D) Regions of ISWI showing significant differential deuterium uptake in the presence of nucleosome than “resting-state” at 60 min of labeling are mapped on to the AlphaFold structure.

### Influence of HSS domain on the rest of the protein

In the structural studies carried so far, variants of ISWI lacking HSS domain have been used. We performed HDX-MS analysis of the ISWI-deletion variant (referred to as the ATPase domain onwards) (Fig. 1A, B, 2C) lacking the HSS domain and the N-terminal 25 residues (Fig. 1A, 2C, S5A) to find whether HSS domain modulates the dynamics of ATPase domain. The ATPase domain has two Rec-A like domains (core-1 and core-2), a feature conserved in all ATP-dependent chromatin remodeling factors and various other protein families (46). Structural mapping of time-dependent deuteration shows that the ATPase domain, like the FL-ISWI, also samples multiple intrinsic conformations (Fig. 2C, S6). We did not observe any region with significant increase in deuteration of the ATPase domain in comparison to the corresponding region in FL-ISWI, suggesting that deletion of the HSS did not deprotect the ATPase domain (Fig. S5B). We did not obtain exactly same peptides for the ATPase-domain in FL-ISWI and deletion variant, therefore we binned the fractional uptake values for 50 residue non-overlapping bins and compared the deuteration between corresponding regions from FL-ISWI and deletion variant. Our analysis suggests that HSS domain does not strongly interacting with the ATPase domain. However, previously reported cross-linking coupled mass spectrometry (XL-MS) experiments captured some interactions between ATPase domain and HSS domain (12). From XL-MS study and our data the HSS domain appear in a very dynamic state and probably transiently interacts with the ATPase-domain or indicating that ISWI exists in multiple conformations with HSS domain being either closer or away from the ATPase domain.

### Conformational changes upon ATP binding

Nucleosome sliding by ISWI involves concomitant steps of ATP and nucleosome binding. To dissect the conformations adopted by ISWI during these individual binding events, we first monitored the conformational dynamics of ISWI in the presence of the non-hydrolysable ATP analogue, adenylyl-imidodiphosphate (AMPPNP) (Fig. S7A). To analyze the effect of ATP binding on the conformational dynamics of ISWI, the deuterium uptake of apo-ISWI was subtracted from that measured in the presence of excess of AMPPNP, and the difference was plotted across the ISWI sequence (Fig. S8B). We observed significant changes in the deuterium uptake in four major regions. The N-terminal region (from 11 to 35) showed increased deuteration (Fig. S8C, D, S7). This region forms a part of the ppHSA domain, which has been shown to strengthen the interaction of the NTR with core-2 of the ATPase domain (13). Different motifs (regions) of ISWI have been shown to be involved in ATP-hydrolysis (active site formation) (17, 47, 48). The region from residues 152 to 163, encompassing motif-I, shows a decrease in deuterium uptake at shorter labeling times. The movement of motifs-I, -II and -VI closer to each other has been reported to occur during ATP-hydrolysis (17, 47, 48). The proximity of these motifs might decrease their conformational fluctuations and lead to a decrease in deuterium uptake. Residue stretch 167-191, shows an increase in deuterium uptake (Fig. S8C, D, S7). XL-MS experiments have shown that Tyr169 and His172 in this region are involved in interaction with Met578 of the core-2 domain (35). From our data, it appears that upon AMPPNP binding, the region from 152 to 191 (covered by peptides 152-163 and 169-172) undergoes a conformational reorientation leading to alignment of motif-I with other motifs involved in ATP-binding and perturbation of inter-core interactions involving residues Tyr169 and His172. Establishment of interactions towards the N-terminus and disruption of interactions at the C-terminus of this region might lead to the dampening of fluctuations at its N-terminus and amplification of fluctuations at the C-terminus, as observed. Another region with decreased deuterium uptake in comparison to that measured in ISWI alone spans from residues 329 to 340 (Fig. 8SC, D, S7). Two of the residues, Lys337 and Pro338, in this region have been shown to be involved in interdomain interactions, and the mutation of Lys337 modulates the ATPase activity (49). We observe modulation of conformational dynamics in different regions of ISWI, some of which are implicated in ATP binding/hydrolysis or inter-domain interactions. These data suggest that binding of ATP induces active site formation and decrease in some interdomain interactions.

### Nucleosome binding induces changes in ATPase core and HSS domains

Although conformations of different domains of ISWI in the absence or presence of nucleosome/ATP-analogues (17,21,50) have been studied, but the conformation attained by FL-ISWI bound to nucleosome have not been studied. To find the conformation adopted by FL-ISWI upon binding to a nucleosome (Fig. 3A), the difference in deuterium uptake between the nucleosome bound state and unbound resting-state (ISWI alone) of ISWI was plotted across the ISWI sequence (Fig.3B). Several regions from residue 793 to 834 show a decrease in deuterium uptake in ISWI-nucleosome complex as compared to ISWI alone (Fig. 3B, C, S9). These residues fall within the SANT region of the HSS domain of ISWI and the region from 793 to 834 contains several conserved hydrophobic and polar residues (51). Similarly, region 764-783 belonging to HAND regions containing three conserved residues, L781, T782 and E783 also shows decreased deuterium uptake (Fig. S9). Binding of HSS domain to extra-nucleosomal DNA or histone H3 tail (15, 51) can dampen the intrinsic dynamics of these regions, resulting in decreased deuterium uptake. Another region in the NTR (residues 24-29 and 33-41) shows decrease in deuterium uptake, implying a dampening of conformational fluctuations of this region upon binding to nucleosome (Fig. 3B, C, D, S9). An alpha-helical region of core-1 from residues 329 to 340 shows increase in deuterium uptake in ISWI-nucleosome complex (Fig. 3B, C, D, S9). Residue 338 in this region has been shown to be involved in the core-1 to core-2 interaction, while Lys337 has been shown to be involved in the interaction between core-1 and the N-terminus (49). Conformational change of this region to a more open-state can perturb the interactions between core-1 and core-2, facilitating the swinging-over of core-2 for nucleosome binding. The alpha-helical region of core-2 from residues 478 to 493 shows a decrease in deuterium uptake, implying that this region binds to the DNA as also shown by previous studies (21). Surprisingly, we did not observe any change in NegC region, possibly because it is highly dynamic in the resting-state conformation, again indicating that the NegC interaction with ATPase core is transient. From our data, it appears that the HSS domain binds to the extra-nucleosomal DNA/H3 tail. The ATPase core undergoes conformational change which seems to result in amplification of dynamics at the interface of core-1 and core-2 and protection of helix from core-2 which binds to nucleosomal DNA (Fig. 3B, D). Interestingly, we found that the regions of ISWI which undergo conformational change upon binding to nucleosome, fluctuate between “open” (high-deuterated) and “closed” (low-deuterated) conformations in the resting-state (Fig. 3D, 2A). Sampling of “open” conformation transiently can facilitate ISWI-nucleosome interactions.

### ATPase core attains different conformations in the nucleosome-bound and nucleosome-sliding states

After binding to the nucleosome ISWI has to transition to the sliding mode. However, determining the conformational dynamics of the FL-ISWI or other chromatin remodelers during actual nucleosome sliding step has proven to be challenging and has not been achieved. To address this problem, we monitored the conformational dynamics of FL-ISWI during nucleosome sliding by HDX-MS. We reconstituted the *in vitro* nucleosome sliding system which has been used extensively to study nucleosome sliding by ATP-dependent chromatin remodeling factors (4,11,22,51). In this assay, ISWI slides a centrally positioned mono-nucleosome towards the end of the DNA template (Fig. 4A, S11A). ISWI, mono-nucleosome and ATP were mixed and D_2_O labelled. After different incubation times, samples were quenched and analyzed by mass spectrometry (Fig.4B). To specifically dissect the conformational changes that ISWI undergoes while switching from a nucleosome-bound state to sliding state, we analyzed the difference in deuterium uptake of ISWI between the two states. A significant change in the deuterium uptake of ISWI during nucleosome sliding relative to that of nucleosome-bound state was observed in several regions from the NTR, core-1, core-2, NegC as well as HSS domains (Fig.4C). Mapping of these regions on the structure of ISWI shows that ISWI undergoes dramatic conformational changes upon transitioning from nucleosome-bound to nucleosome-sliding state (Fig. 5A). We also mapped the regions undergoing conformational change on a deletion variant of *C. thermophilum* ISWI (CtISWI) in complex with nucleosome (21) to determine the spatial arrangement of different regions of ISWI with respect to the nucleosome (Fig. 5B).

**Figure 4.**
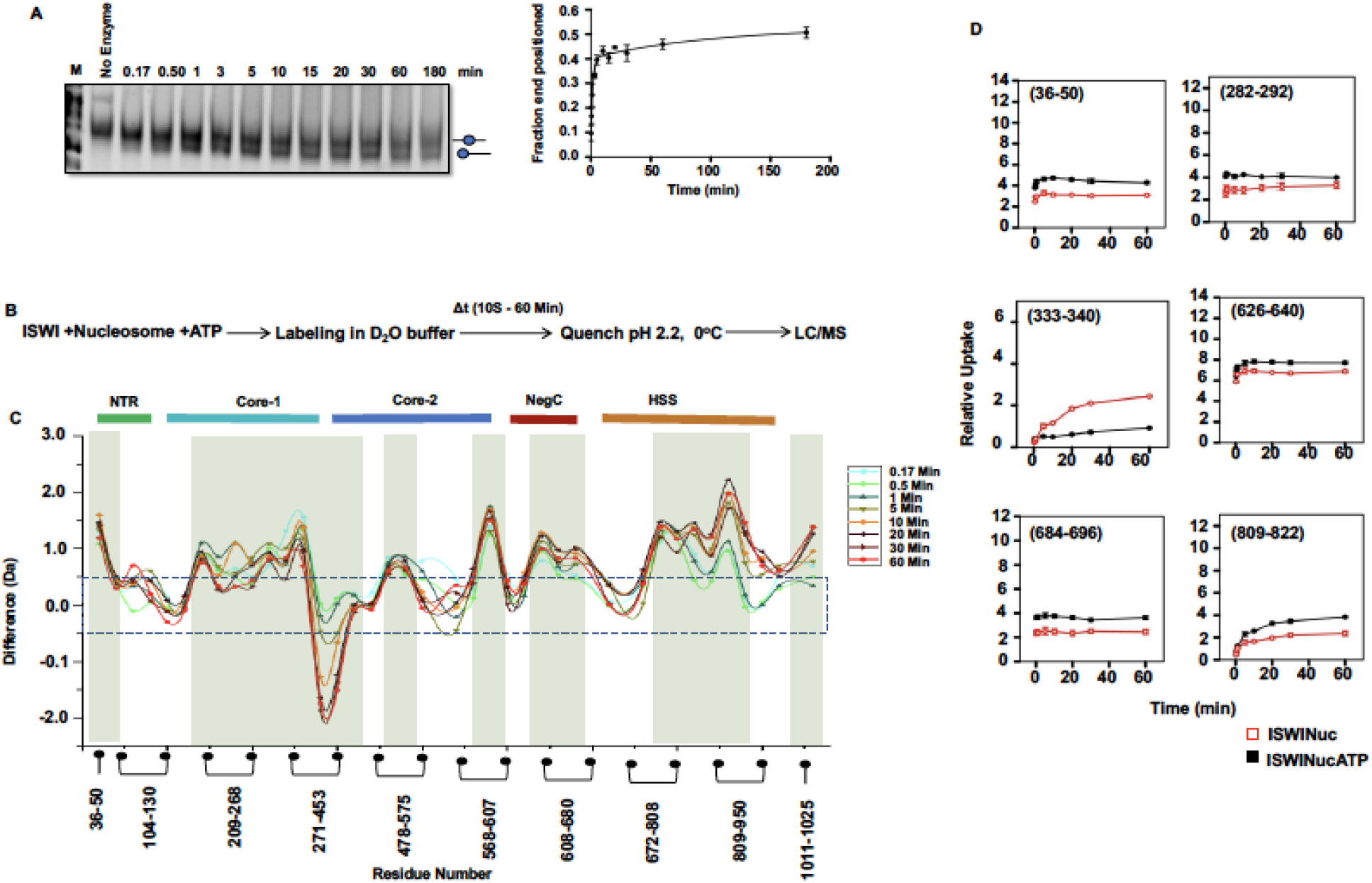
Conformational dynamics of ISWI during nucleosome sliding: (A) Representative native-PAGE showing the increase in the intensity of the band corresponding to end-positioned nucleosome (lower band) and the quantitation of fraction end positioned nucleosome with time. (B) Flow chart showing the main steps of the HDX-MS experiment for the nucleosome sliding. (C)Difference (subtraction) plot of ISWI during nucleosome sliding and ISWI-nucleosome bound state at different labelling times. Differences in deuterium exchange more than ±0.5 Da are considered significant (dashed box). Shaded regions show regions with significant change. (D) Representative deuterium uptake plots of different regions of ISWI during nucleosome sliding (black) or in nucleosome-bound state (red) obtained from three different sliding experiments and two different nucleosome binding experiments, respectively. Error bars show the standard deviations.

**Figure 5.**
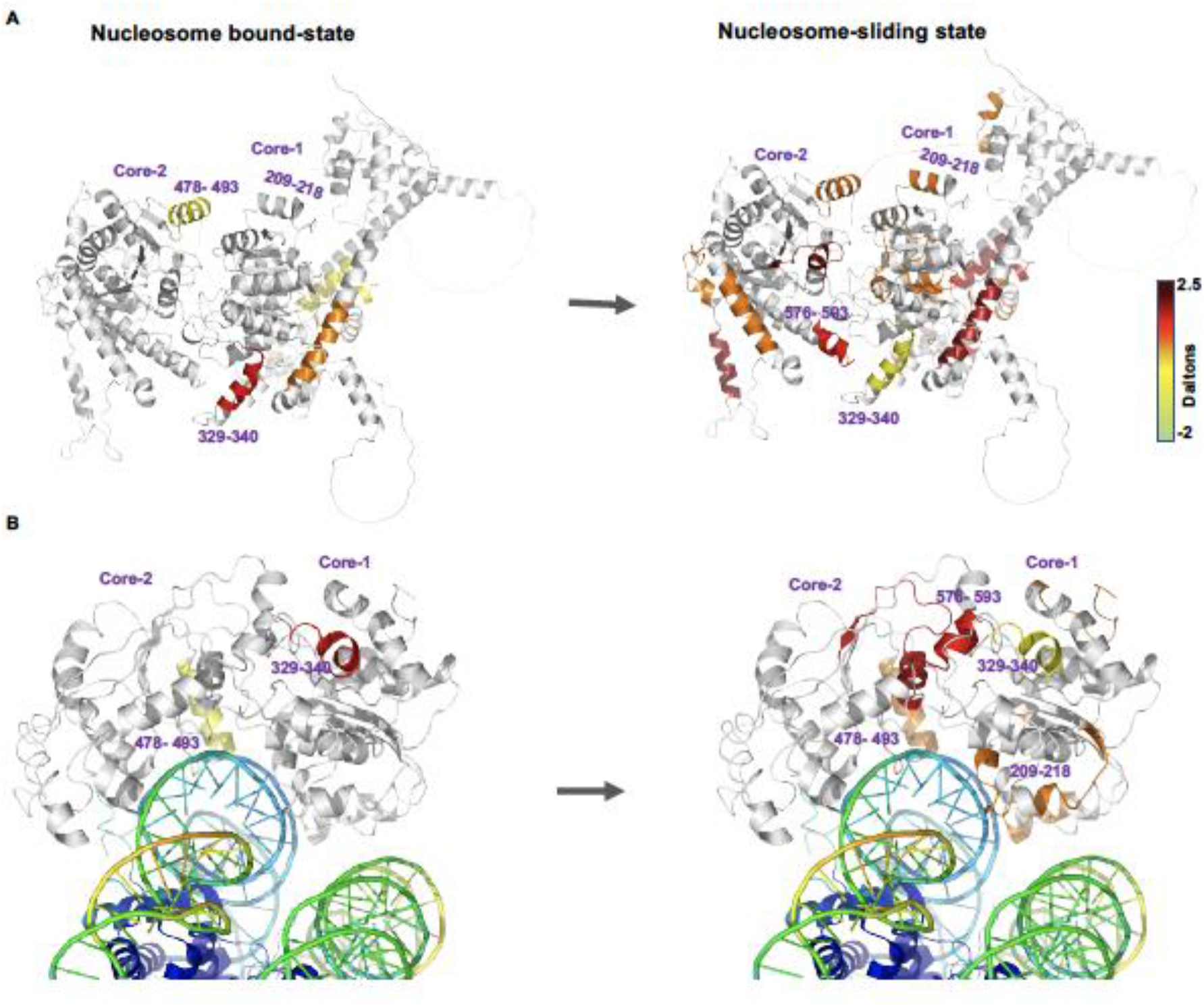
Conformational transition of ISWI from the nucleosome-bound to nucleosome sliding state: (A) Regions of ISWI undergoing conformational change upon binding to nucleosome (relative to ISWI alone) and during nucleosome sliding (relative to bound-state) mapped on to the FL-ISWI alpha-fold structure (ID:AF-Q24368-F1). (B) Regions of the ATPase domain undergoing conformational change in the ISWI-nucleosome complex mapped on to the CtISWI ATPase domain-nucleosome complex (PDB code 6PWF) (left) and regions undergoing conformational change during sliding (relative to bound-state) (right).

From the N-terminal region, residues from 36 to 50 show more deuterium uptake during sliding than the corresponding residues in nucleosome-bound state. This region forms a part of major helix and contains two highly conserved arginine residues and a leucine residue (Fig. S10). From core-1, four regions 115-130, 209-218, 256 to 268 and 271-281 show an increase in deuterium uptake (Fig.4C, D, Fig. 5, 6 Fig. S11, 12, 13) during nucleosome sliding. Two of these regions 209-218 and 256-268 mapped on the CtISWI-nucleosome complex are involved in binding to the nucleosomal DNA. Sequence segment 209 -218 corresponds to the sequence segment 266 -273 of CtISWI which contains a highly conserved arginine residue involved in binding to the phosphate backbone. The region from residues 256 to 268 contains the highly conserved His259, Arg260 and Lys262 which are also involved in binding to the phosphate backbone (Fig. 5B, S10). However, the region of core-1 from residues 329 to 340, mapping away from the nucleosome binding site shows a decrease in deuterium uptake during the sliding in comparison to the nucleosome bound state. This suggests that this region gets exposed to the solvent in the nucleosome bound state but seems to switch to a buried conformation during nucleosome sliding. From core-2 three regions: 478 to 493, 536 to 552 and 576 to 593 show an increase in deuterium uptake during nucleosome sliding than that of nucleosome-bound state (Fig. 4C, 6). The region spanning residues 478 to 493 constitutes the alpha-helix which binds to the DNA *via* the highly conserved His483 and Arg486 (Fig. 5B). Interestingly this region attains a buried conformation in the nucleosome bound-state as it remains bound to the phosphate backbone but to allow translocation of DNA this motif has to move and hence, shows an increase in deuterium exchange. The regions spanning residues 536 to 552 and residues 576 to 593 (Fig. 4C, Fig. 6) are away from the nucleosome binding site but are at the interface of the core-1 and core-2 of the ATPase domain (Fig. 5B). Met578, in the region spanning residues 576 to 593, seems to act as a hub for interdomain interactions (35). It is involved in the core-1 to core-2 interaction, as well as in the interaction with Trp942 of the SLIDE region of the HSS domain. This region is also in close proximity to the region 329-340 of core-1 which also shows increase in deuterium uptake. We suggest that movement of the two core domains during nucleosome sliding occurs around this site possibly to adjust the positions of other motifs involved in DNA translocation.

**Figure 6.**
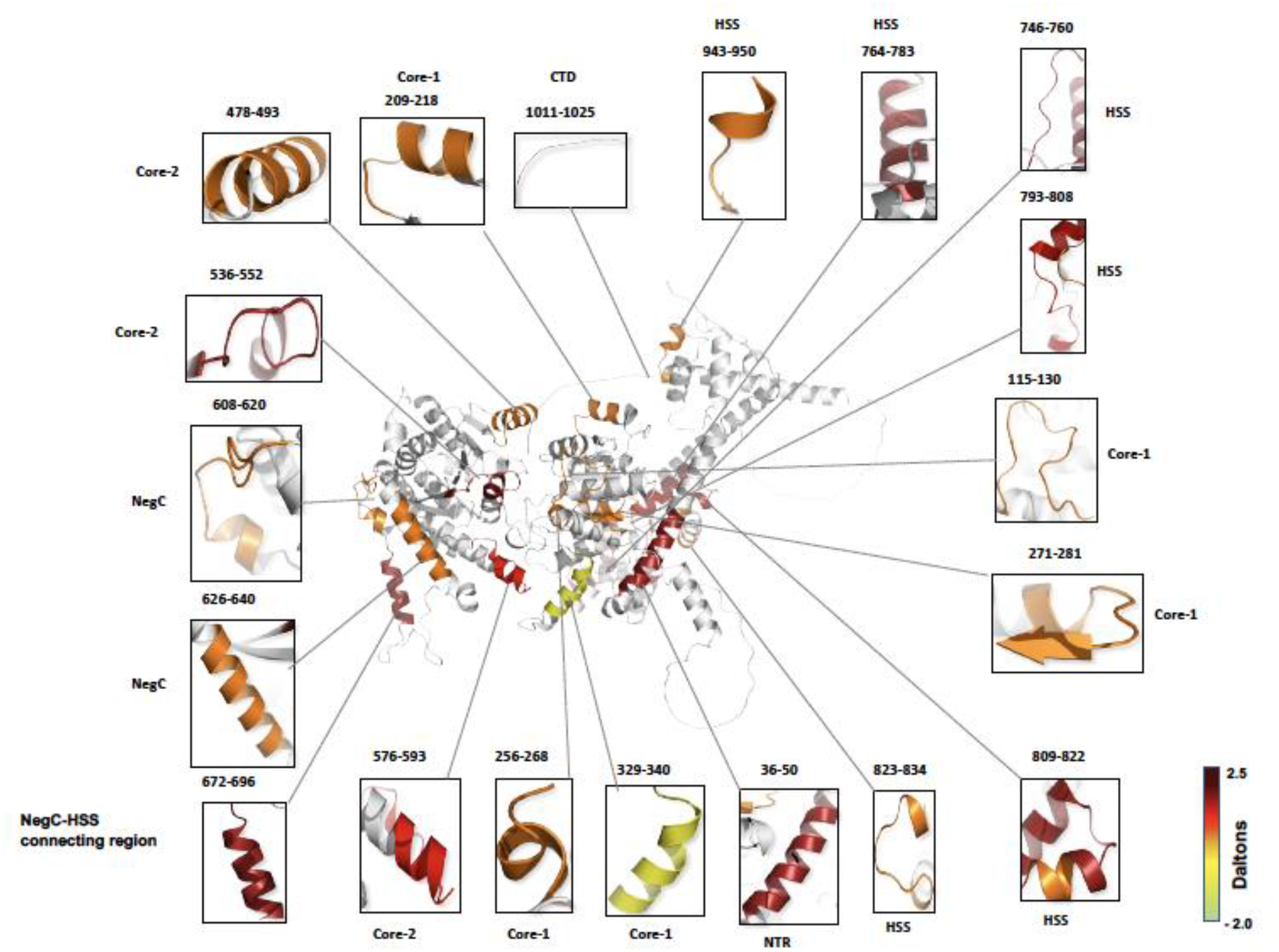
Regions of ISWI undergoing conformational change during nucleosome sliding: Regions of ISWI showing significant differential deuterium uptake during nucleosome sliding relative to the nucleosome-bound state at 60 min of labeling are mapped on to the ISWI Alpha fold structure. The amino acid sequence of these regions is shown in figure S10.

In summary, we have identified eight regions of the ATPase domain from nucleosome binding sites as well as away from the nucleosome binding sites, which undergo conformational change during nucleosome sliding. The increase in deuterium uptake of regions during sliding, which bind to nucleosomal DNA in the ISWI-nucleosome complex, suggests that the interactions between these regions and the nucleosome are broken to allow translocation of DNA. From these data it appears that during nucleosome sliding, the NTR of ISWI becomes more exposed to solvent possibly by moving away from the ATPase-core, and that two lobes of the ATPase domain undergo conformational reorientation which is different than the conformational change observed upon nucleosome binding. Hence, the conformation attained by ATPase domain upon binding to the nucleosome is very different from the conformation attained by it during sliding.

### C-terminal domains undergo major conformational change during nucleosome sliding

The conformations attained by Neg-C and HSS domains during nucleosome sliding remain completely obscure. Here, we were able to monitor the conformational changes in these two domains as well as the region at the end of the C-terminus. From the Neg-C region, residues 608- 620 and 626-640, show an increase in deuterium uptake during nucleosome sliding (Fig. 4C, 6, S11, S12, S13). These regions contain several highly conserved polar residues (12). Three such residues (DED) 619-621 are highly conserved across species (Fig. S10), implying their possible role in ISWI function. The stretch from residues 672 to 696 falling in the region of ISWI connecting NegC and HSS domain, containing highly conserved KRERK motif (Fig. S10) also shows increase in deuterium uptake (Fig.6). The conformational dynamics of this region might be involved in communication between HSS and NegC during nucleosome sliding.

The HSS domain shows increased deuterium uptake in four regions: 746-760, 764-783, 793-808 and 943-950 (Fig. 4C, 6). The residues from 746 to 760 belong to helix-2 of the HAND domain, having a conserved lysine residue at position 748. Residue stretch 793-808 is part of an alpha-helix of SANT regions (Fig. 6) containing several conserved residues (51) as well as Lys810 (Fig. S10) that is involved in interaction with the core-2 domain (35). Interestingly, the region 764- 783 shows a decrease in deuterium uptake upon nucleosome binding, but during nucleosome sliding this region shows increase in deuterium uptake. This observation suggests that this region switches from being in a bound-state (in ISWI-nucleosome) to a dynamic state during nucleosome sliding. Surprisingly, we also found an increase in deuterium uptake of C-terminal domain, 1011- 1025 (Fig. 4C, 6), although it has been predicted to be in an unstructured form. Therefore, during nucleosome sliding, a) Neg-C region seems to move away from ATPase-core resulting in increase in its deuterium uptake b) the dynamics of HSS domain increases which will be required to constantly sense the length of extra-nucleosomal DNA and coupling with ATPase activity and c) the region connecting Neg-C and HSS domain also shows enhanced fluctuations. Taken together, we have for the first time comprehensively identified eighteen regions of ISWI from the NTR, ATPase domain, Neg-C, HSS domain and CTD, which undergo conformational change when ISWI switches from nucleosome-bound to nucleosome-sliding state. We also analyzed the difference between deuterium uptake of ISWI during nucleosome sliding and ISWI alone, and from this analysis 13 regions from different domains of ISWI were identified which show conformational change during nucleosome sliding (Fig. S14, S15, S16).

### Relationship between intrinsic and sliding conformations of ISWI

Intrinsic dynamics of enzymes have been shown to be linked with the conformational changes involved in their catalytic activity or folding/unfolding reaction (29,37, 52). We examined whether there exists any relationship between the intrinsic conformational heterogeneity of ISWI and conformations adopted by ISWI during nucleosome binding or sliding. Analysis of conformations sampled intrinsically by ISWI in the resting-state and during nucleosome sliding, at the same labelling times, show a good correlation, in terms of deuterium exchange and the regions involved in conformational dynamics (Fig. 7A, B, C, D, S17). Surprisingly, all the regions undergoing conformational change during nucleosome sliding, in comparison to nucleosome-bound or resting states, fall in the Class-b and Class-c regions (Fig. 7B, C, E, S16), both of which fluctuate intrinsically in resting-state. These analyses show that there is some level of similarity between conformations sampled intrinsically by ISWI with those sampled by ISWI during nucleosome sliding. Therefore, we suggest that conformational changes that ISWI undergoes during nucleosome sliding can be initiated from its intrinsic ensemble of interconvertible conformations. Furthermore, regions of ISWI showing altered deuterium uptake upon binding to AMPPNP or nucleosome also fall in the Class-c regions, substantiating the observation that conformational changes required for the binding of ATP or nucleosomes can be initiated from the already existing intrinsic conformational fluctuations of ISWI. Intrinsic conformational dynamics of ISWI similar to that observed during nucleosome binding or sliding suggests that these intrinsic fluctuations are not random but have possibly evolved to achieve the specific activity of ISWI.

**Figure 7.**
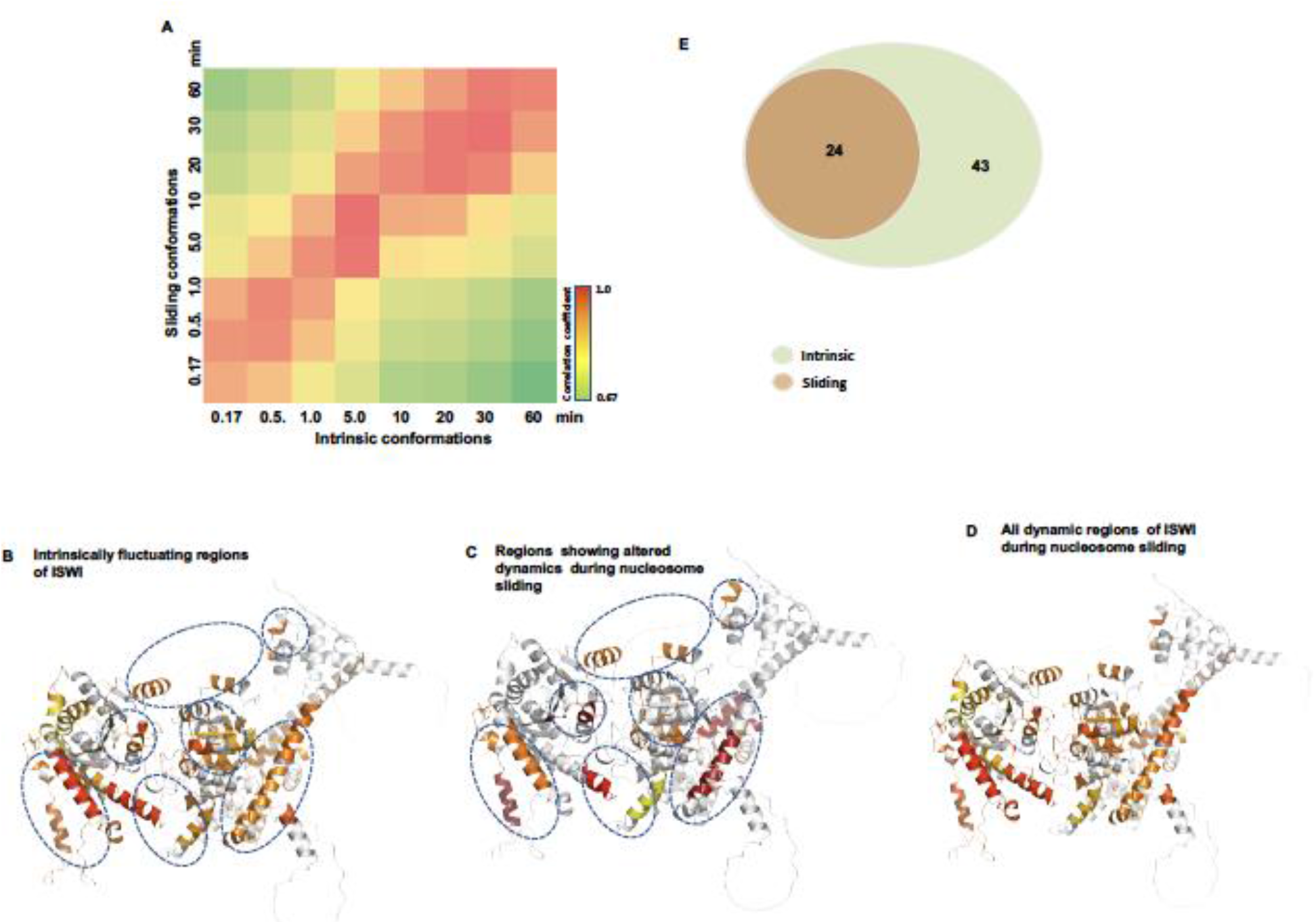
Relationship between the intrinsic and sliding conformations of ISWI: (A) Heatmap showing the correlation between intrinsic and sliding conformations of ISWI. Conformation sampled by ISWI intrinsically (A) and during nucleosome sliding at 10 min of labelling (D). (C) 3D structure showing regions with increased deuterium uptake than corresponding in nucleosome-bound state. Dashed ovals on the 3D structure represent corresponding regions fluctuating intrinsically as in panel A. (E) Overlap between regions (peptides) showing increased deuterium uptake during nucleosome sliding (in comparison to nucleosome-bound state) and intrinsically fluctuating regions from Class-b and Class-c.

## Discussion

In this study, the conformational dynamics of FL-ISWI was monitored for the first time at different stages of the nucleosome sliding process as well as in the resting-state. Given the structural complexity of ISWI, it has proven challenging to study conformational transitions of FL-ISWI during nucleosome sliding. To overcome this problem, we applied in solution based method, HDX-MS to uncover the conformations attained by ISWI across entire sliding process. Our data provides insights into two important mechanistic questions: a) How does ISWI switch from the “auto-inhibited” state to the nucleosome-bound state? and b) What are the conformational changes ISWI undergoes to transition from the nucleosome-bound state to the nucleosome-sliding state? The structure of FL-ISWI from any organism has not been solved. From biochemical, mutational, cross-linking and structural studies (with deletion variants), ISWI has been proposed to attain an auto-inhibited state in which N-and C-terminal regions interact with the ATPase domain attaining a “closed” conformation (11,12,17). To bind the nucleosome, ISWI must overcome this auto-inhibition and attain a conformation competent for nucleosome binding. Indirect evidence from kinetics of ATP hydrolysis suggests that ISWI can exist in two sub-populations which differ in their ATPase activities (20). Also, the structure of nucleosome-bound-ATPase core from two different species indicate different conformations of the NegC (17,53). Moreover, it has proven difficult to fit all the observed interdomain cross-links of ISWI in a unique conformation (12). These reports point towards the possibility that there exits some intrinsic conformational heterogeneity in ISWI. The intrinsic conformational heterogeneity of proteins has been linked with catalysis of enzymes, binding of ligands and protein-protein interactions (27, 29,52, 54-58). It has proven challenging to detect conformational sub-states of proteins and to link them with their designated function. The conformations which significantly differ from the most stable state get populated to very low levels because of large energy difference between them and the most stable state (Boltzmann-distributed), making them difficult to detect. However conformational sub-states which get populated significantly are very similar to the most stable, again making their detection difficult. Here, we provide direct evidence that FL-ISWI as well as its ATPase domain sample an ensemble of conformations in the resting-state. Conformations showing more deuterium uptake are “open” and conformations taking up less deuterium are relatively “closed”. Several of the regions fluctuating intrinsically between low and high deuterated states are involved in auto-inhibition and inter-domain interactions. Therefore, “closed” conformations may be more auto-inhibited, and “open” conformations less auto-inhibited or competent for nucleosome binding (Fig. 2B). Under native conditions “closed”/folded conformation of ISWI will be most stable and therefore most populated while as “open” conformation will be unstable and least populated (Boltzmann-distributed) (Fig. 8A). Although both conformations exist together but lower fraction of molecules in the “open” state makes them undetectable by conventional methods. However, shuttling of ISWI molecules between “closed” and “open” conformations makes them detectable by continuous labelling HX-MS. Observation of unimodal mass spectra shifting towards higher mass with increase in labeling time suggests that the exchange occurs by local unfolding/opening events.

**Figure 8.**
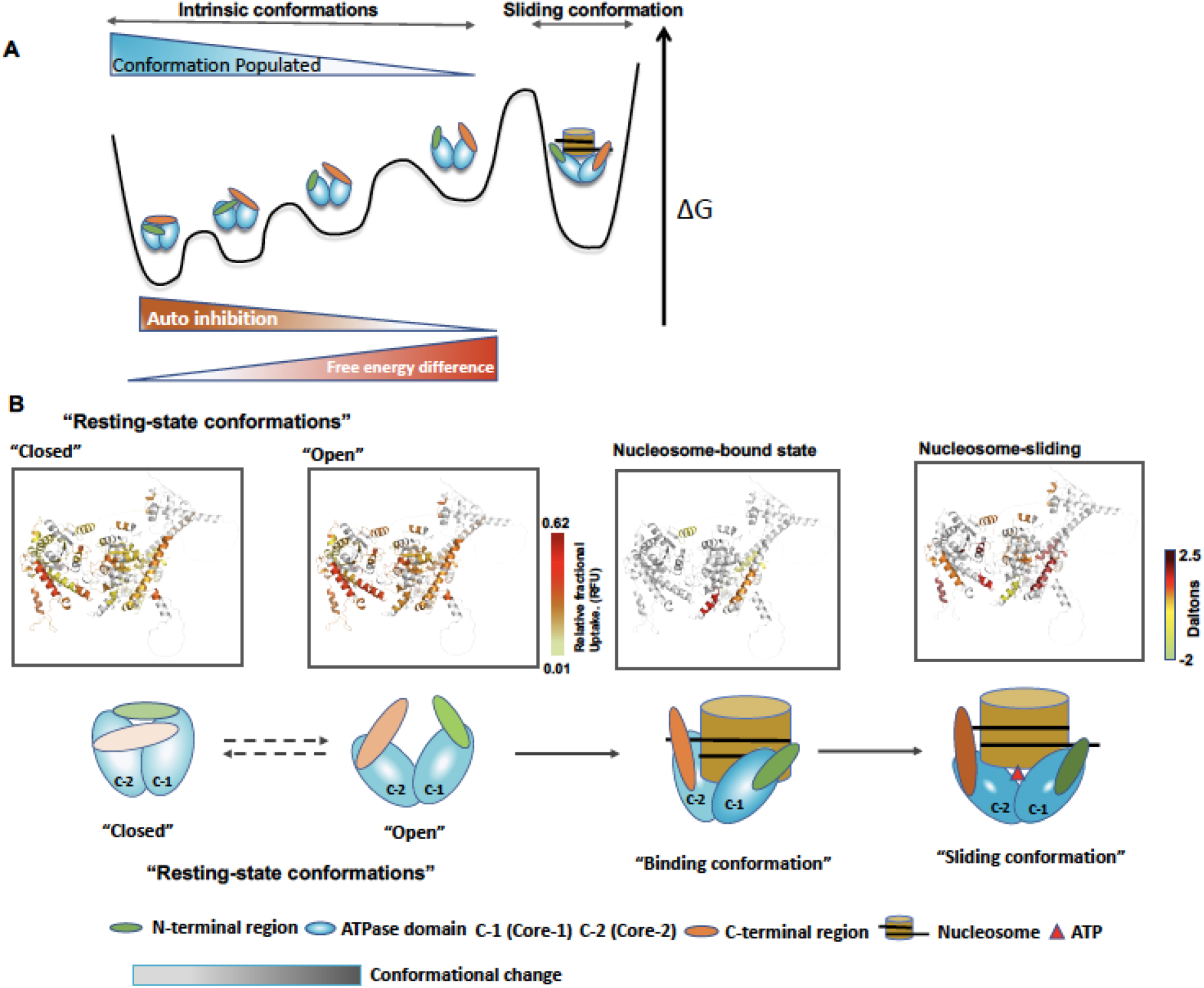
Conformational landscape sampled by ISWI during nucleosome sliding: (A) Schematic showing ISWI samples multiple conformations in resting-state. Different conformations have varying solvent-exposure and hence, show differential deuterium uptake. Conformations having more solvent exposed regions will be more “open”, having less interdomain interactions and less auto-inhibition. But “open” conformations will be sampled less frequently due to a large difference in free energy from the most stable state (closed). These equilibrium conformations can be exploited for sliding. Relative energies and placement of different conformations are drawn for illustration only. (B) Schematic showing intrinsically sampled “closed” and “open” conformations of ISWI. Dashed arrows indicate there can be more partially open conformations between the two states. “Open” conformation will favor nucleosome binding and shift the equilibrium towards bound-state. Upon ATP hydrolysis conformational change across all domains of ISWI occurs to slide the nucleosome. Conformations attained by ISWI with different levels of deuterium uptake in different regions are shown on top. Change in conformation of ISWI in nucleosome-bound state is relative to open-state and change in conformation of sliding-state is relative to nucleosome-bound state. Change in intensity of color in schematic indicates conformational change.

Given the coincident location for binding of auto-inhibitory regions and nucleosomal epitopes on the ATPase domain, in “open” conformation ISWI-nucleosome interactions will be facilitated but avoided in the “closed” state in which intra-ISWI interactions are favored. Shuttling of ISWI between the “open” and “closed” conformations will allow a time window during which ISWI-nucleosome interactions will be favored, as proposed previously also (19). Therefore, binding of nucleosome to the “open” state will shift the equilibrium towards bound state and lead to initiation of nucleosome sliding (Fig. 8B). Based on these observations we propose that ISWI is “auto-regulated” instead of “auto-inhibited”. Cells may regulate the activity of ISWI by modulating its dynamics between “closed” and “open” states.

In agreement with previous structural studies (21, 59) we also identified the structural elements of the ATPase domain which interact with the nucleosome in an ISWI-nucleosome complex. We also observed a conformational change of a region (329-340) located at the interface of core-1 and core-2 when ISWI transitions from the “resting-state” to the nucleosome-bound state. This conformational change resulted in exposure of this region to the solvent possibly due to swinging over of core-2 domain upon binding to the nucleosome (21, 59, 60). The HSS domain attains a solvent-protected state in nucleosome bound-state as it shows a decrease in deuterium exchange, in agreement with its role in binding to the extra-nucleosomal DNA (51). Surprisingly, upon binding to nucleosome, no significant change in deuteration of NegC region was observed demanding further investigation into the mechanism by which NegC regulates ISWI activity.

Although structures of ISWI deletion-variants bound to nucleosome have been studied but it has proven challenging to monitor the conformational dynamics of a chromatin remodeler in real-time during nucleosome sliding by a high-resolution/sensitivity probe. Several conformational changes have been predicted to occur in a chromatin remodeler while nucleosome sliding. Cycling of core-1 and core-2 domains of the ATPase domain between states in which they are closer or away from each other has been proposed to be involved in nucleosome sliding (59). Our data shows that nearly entire ISWI is dynamic during nucleosome sliding and undergoes major conformational changes at regions involved in auto-inhibition, ATPase domain, HSS domain and CTD in comparison to its nucleosome-bound state or resting state. We observe an increase in the dynamics of two regions of core-2 at the interface of the core-1 and core-2 domains. We observe an increase in deuterium uptake at the regions of ATPase domain during sliding which have been shown to bind the nucleosomal DNA in ISWI-nucleosome complex, suggesting that these interactions are broken when ISWI transitions from the nucleosome-bound to nucleosome-sliding state to allow movement of DNA towards the exit-site. However, the region of core-1 which got exposed to solvent in the nucleosome-bound state switched to a buried conformation in the nucleosome-sliding state, demonstrating further the complexity of conformational changes ISWI undergoes during nucleosome sliding.

Conformational coupling between HSS and the ATPase core has been proposed for coordinated ATPase and sliding activities of ISWI in order to regulate the nucleosome spacing. But the conformations attained by NegC and the HSS domain during nucleosome sliding have remained obscure. We show that NegC and HSS domain undergo conformational changes during nucleosome sliding relative to the nucleosome-bound state. Conformational change in NegC will be crucial for activation of ISWI as it negatively regulates the ATPase core. The region connecting NegC and the HSS domain also undergoes conformational change during nucleosome sliding, as it may be involved in communication between the HSS domain and NegC. HSS domain functions as flanking DNA length sensor and hence, is expected to constantly function in coordination with the ATPase domain to regulate the nucleosome sliding. Observation of increased deuterium uptake in the HSS domain during nucleosome sliding suggests that the HSS domain possesses enhanced dynamics adopting different conformations as DNA is being translocated. This contrasts with decrease in deuterium uptake of the HSS domain observed upon binding to the nucleosome, where it just remains bound to the extra-nucleosomal DNA.

In summary, we have precisely identified the structural elements across FL-ISWI, which undergo conformational change during nucleosome sliding. However, there are some regions (most of them unstructured) for which coverage was not observed in mass spectrometry, and it is possible that some of those regions are also involved in binding or nucleosome sliding.

Based on previous studies and our data, we propose a conformational cycle through which ISWI goes to slide a nucleosome (Fig. 8): i) ISWI intrinsically fluctuates between “closed” and “open” conformations ii) the “open” conformation favors nucleosome binding, leading to dislodging of the NTR and movement of the ATPase core domains to achieve the nucleosome-bound state iii) the NegC region gets displaced possibly due to the binding of HSS to linker DNA and/or ATP hydrolysis iv) the ATPase domain undergoes conformational change due to breaking of contacts established with the nucleosome during binding and increased dynamics away from the nucleosome binding site possibly to translocate DNA v) the HSS domain switches from being just bound to extra-nucleosomal DNA to a dynamic state sensing the length of linker DNA which is coupled to nucleosome sliding activity of the ATPase motor. The precise temporal order in which these events occur demand further investigation.

We suggest that the intrinsic conformational dynamics of ISWI between “closed” and relatively “open” conformations can establish an “auto-regulation” mechanism of ISWI activity. This study has broad implications for understanding the mechanics of not only ATP-dependent chromatin remodeling factors of ISWI family across species, but also of several other protein families containing Rec-A like domains (core-1 and core-2) (40). Like ISWI, other Rec-A like proteins might also possess intrinsic dynamics which can play role in their function.

## Acknowledgments

We thank Prof. Felix-Plantz Muller for providing ISWI constructs and Prof. Geeta Narlikar for providing histone constructs. We also thank Prof. Felix-Plantz, Prof. Robert Kingston, Prof. Nicole Francis, and our group members for their valuable suggestions and discussions on the manuscript.

## Supporting Information

This article contains supporting information.

