## supplemental material for "HDX-MS reveals concealed conformations of ISWI during different stages of nucleosome sliding"

**Supporting Information**

**List of items included**

1. Supplementary Text
2. Supplementary table
3. Supplementary figures and legends

**Materials and Methods:**

**Expression and purification of ISWI and its mutant**

pPROEX-HTb–based expression plasmids with genes encoding *Drosophila* ISWI full length and ISWI 26–648 (truncated version, ATPase domain) were a kind gift from Prof. Felix Mueller-Planitz (TU Dresden, Germany). The plasmids were transformed into BL21-Gold (DE3) competent cells. The expression and purification of both the proteins of ISWI were performed as mentioned earlier (16). Briefly, the cells with recombinant plasmids were grown at 37$^{o}$C in shaking incubator in LB broth (Merck) supplemented with 100 µg/µl ampicillin (Sigma Aldrich) and induced at OD of 0.8 at 600 nm, with 0.2 mM IPTG (Sigma-Aldrich) at 20 $^{o}$C overnight. Cells were harvested by centrifugation and resuspended in cold lysis buffer (10ml/L of culture) 15 mM Tris-chloride pH 8, 300 mM NaCl, 10% glycerol, 0.2% triton X, 10 mM imidazole,500 ug/ml lysozyme and protease inhibitors (cOmplete, EDTA-free, Roche). Cells were lysed by sonication (amplitude 60%, 10 second on/off pulse for 10minutes). Lysate was cleared by centrifugation at 16000 rpm for 30 min using JA 25.50 rotor (Beckman Coulter) at 4$^{o}$C. The lysate was applied to Ni-NTA beads (Qiagen) pre-equillibrated with buffer A (15 mM Tris-chloride pH 8, 500 mM NaCl, 10% glycerol, 0.05% tween 20 and 10 mM imidazole). The mixture was incubated for three hours and then applied on a Econopac column (Biorad) and the flow through was collected. The beads were then washed 10 column volumes (CV) of buffer A and 5 CV of buffer A supplemented with 30mM imidazole. Elution was performed using buffer A with 250-300 mM imidazole. The purity of the collected fractions was analyzed on 10% and 8% SDS-PAGE for ISWI 26–648 and FL-ISWI, respectively. The partially purified protein fractions were pooled and incubated overnight with TEV protease (10:1 mass/mass ratio) (62) for tag removal in buffer B (15mM Tris-Cl pH 7.4, and 1 mM DTT). Uncleaved protein and TEV was removed by applying the mixture to Ni-NTA beads (Qiagen) equilibrated with buffer B. The flow through was collected and loaded on to monoS cation exchange column (GE Healthcare) preequilibrated with buffer C (15 mM Tris-chloride pH 7.4, 50 mM NaCl, 10% glycerol) connected to Akta (GE). The proteins were eluted by 0.05-1 M NaCl gradient and different fractions were collected separately. The purity of factions was analyzed by SDS-PAGE and the desired fractions were pooled and applied to a gel filtration column, Sephacryl S200 16/60, (GE Healthcare) preequilibrated with 20 mM HEPES-KOH pH 7.6, 200 mM potassium chloride, 0.2 mM EDTA, 5 mM dithiothreitol. The pure fractions were pooled, concentrated, flash frozen and stored at -80$^{o}$C. The concentration of purified ISWI and ATPase domain was determined by measuring absorbance at 280 nm using, extinction coefficient(ε) of 119950 $cm^{-1}$ $M^{-1}$(43), and 75860 $cm^{-1}$ $M^{-1}$. Extinction coefficients of ATPase domain was obtained by using ExpasyProtParam tool. The concentration of the purified proteins was also checked by SDS-PAGE using BSA of known concentrations as standard.

**Expression and Purification of Histones**

The Xenopus laevis core histones were purified as earlier (2) with minor modifications. Histone constructs were kind gift from Prof. Geeta Narlikar (UCSF) and Dr Altaf Bhat (CIRI, University of Kashmir). Briefly, the histones were expressed as inclusion bodies in Bl21-DE3 cells. The inclusion bodies were dissolved in denaturing buffer (6 M guanidinium hydrochloride solution, containing 20 mM sodium acetate (pH 5.2) and 1 mM DTT). The solubilized inclusion bodies were desalted against low salt buffer (7 M urea, 10 mM Tris (pH 8), 200 mM NaCl, and 2 mM 2-mercaptoethanol). The histones were purified through anion (DEAE-Sigma Aldrich) and cation exchange chromatography (SP-Sepharose, GE-Healthcare) held in tandem. The histones were eluted from cation exchange column using high salt buffer (7 M urea, 10 mM Tris (pH 8), 1M NaCl, and 2 mM 2-mercaptoethanol). The purified proteins were dialyzed against three changes of miliQ water supplemented with 1 mM PMSF (Sigma) and stored as lyophilized powder. Concentration of the purified histones was checked by absorbance at 276 nm with molar extinction coefficients (ε) of 4050 ($cm^{-1}$ $M^{-1}$), 6070 ($cm^{-1}$ $M^{-1}$), 4040 ($cm^{-1}$ $M^{-1}$) and 5040 ($cm^{-1}$ $M^{-1}$) for H2A, H2B, H3 and H4, respectively (2).

**Reconstitution of H2A/H2B dimer and H3/H4 tetramer**

The histone H2A/B dimer and H3/4 tetramer were reconstituted from purified histones as reported previously (2). The lyophilized histones were denatured in unfolding buffer (6 M guanidinium HCl, 20 mM Tris-HCl, pH 7.5, and 5 mM DTT) for an hour at 22$^{o}$C and equimolar ratios of H2A was mixed with H2B and H3 was mixed with H4. The mixtures were dialyzed against three changes of refolding buffer (2 M NaCl, 10 mM Tris-HCl, pH 7.5, 1 mM Na-EDTA, 5 mM BME) at 4$^{o}$C. The H2A/B dimers and H3/4 tetramers were purified through Superdex S200 16/60 column (GE-Healthcare) and the purified fractions were analyzed by 15% SDS-PAGE and stored at 4$^{o}$C. Concentration of histone H2A/H2B dimer and H3/H4 tetramer was calculated from SDS-PAGE using BSA of known concentrations as standard.

**Mononucleosome preparation:** Mononucleosomes were reconstituted using equimolar concentrations of purified H2A/H2B dimer, H3/H4 tetramer and purified 197 bp of 601 DNA sequence in a high salt buffer (2 M KCl, 10 mM Tris-HCl, pH 7.5, 1 mM EDTA, 1 mM DTT). The mixture was dialyzed over 24 hours against the low salt buffer (0.25 M KCl, 10 mM Tris-HCl, pH 7.5, 1 mM EDTA, 1mM DTT) using double peristaltic pump with flow rate of 1.5 ml/min. The solution was further dialyzed for 3 hours against low salt buffer. Mononucleosome preparation was checked on 4.5 % native-PAGE, pre-run for 30 min at 4 $^{o}$C using 0.25X TBE as running buffer. The sample was loaded and the gel was run for 90 min at a constant voltage of 110 V at 4 $^{o}$C. The gel was stained using sybrgold (ThermoFisher Scientific) and photographed using gel documentation system. For sliding reactions the mononucleosome were buffer exchanged with remodeling buffer. The concentration of monucleosome was determined by using native-PAGE with DNA of known concentrations as standard, as described in Zhou, and Narlikar (63).

**Nucleosome sliding assay**

Sliding reactions were performed by mixing 75 nM ISWI with 75 nM mononucleosome in 20 mM HEPES-KOH pH 7.6, 50 mM NaCl, 1mM MgCl2, 1 mM DTT, 0.1 mM EDTA and 10% glycerol. The mixture was kept at 26$^{o}$C for 5 minutes and the reaction was initiated by adding 1.5 mM ATP (Sigma Aldrich). 10 µl aliquots were taken at different times and the reaction was stopped using 200 ng of plasmid DNA, 10 mM ADP (Sigma Aldrich) and 10% glycerol. Samples were run on 4.5% native-PAGE as done for mono-nucleosomes and the gel was stained for 10 min with syber gold (Thermo Fisher Scientific).

**Hydrogen deuterium exchange**

An exchangeable hydrogen gets replaced by solvent deuterium when a protein undergoes local or global unfolding events. The exchange is influenced by H-bonding, secondary and tertiary structure of protein. These factors determine the protection factor of an exchangeable hydrogen. Therefore, for exchange to take, a site has to overcome these factors to attain an exchange competent solvent accessible state. This is explained by LinderStrom equation (Linderstrøm-Lang,1955 *Chem. Soc. Spec.)*

$$\mathrm{NH\ (closed)} \underset{k_{cl}}{\overset{k_{op}}{\rightleftharpoons}} \mathrm{NH\ (open)} \xrightarrow{K_{int}} \mathrm{ND\ (exchange)}$$

The rate constant of intrinsic exchange, $k_{int}$ is base catalyzed when the pH of the solvent is above 3. $k_{int}$ depends upon the pH, the nature of the amino acid residue, its flanking residues, temperature and salt concentration (64). If $k_{op}$ and $k_{cl}$ are independent

of pH, then the observed exchange rate constant, $k_{HX}$ is given by:

$$k_{HX} = \frac{k_{op} * k_{int}}{k_{op} + k_{cl} + k_{int}}$$

Under EX1 condition, where $k_{op} < k_{cl} << k_{int}$, every unfolding event leads to exchange of one or more deuteriums, and the observed exchange rate constant, $k_{HX} = k_{op}$. Under EX2 condition, where $k_{cl} >> k_{int}$, every unfolding event does not result in exchange, and an equilibrium get established between the folded and the exchange-competent unfolded state and $k_{HX} = (k_{op}/k_{cl})*k_{int}$. Under EX2 conditions observed exchange rate increases with increase in pH because kint increases 10 fold with every 1 unit increase in pH, but observed exchange rate under EX1 condition is independent of pH.

To study the intrinsic dynamics of ISWI and ATPase domain, 2.5µl of purified protein mixed with 3.5 µl protonated buffer were diluted 10 fold (54µl) with deuterated labelling buffer (20 mM HEPES pH7.6, 50 mM NaCl, 0.1 mM $MgCl_2$, 0.1 mM EDTA, 1 mM DTT in $D_2O$) for 10 s to 60 minutes at a final ISWI concentration of 830 nM. The labeling reactions were performed at 26 °C followed by quenching using ice cold 60 µl of quench buffer (100 mM glycine pH 2.2 and 25 mM TCEP in $H_2O$) to minimize back-exchange.

To study the effect of ATP binding, 2.5 µl of purified ISWI, 1 µl of non-hydrolysable analog of ATP AMPPNP (adenylyl-imidodiphosphate, from Sigma) and 2.5 µl of protonated buffer were mixed for 1 min at RT and then labeling was performed by diluting 10 fold (54 µl) with deuterated labelling buffer for 10 s to 60 minutes. The concentration of ISWI and AMPPNP was 830 nM and 1.5 mM (in excess) (18, 31, 33), respectively. The labeling reactions were performed at 26 °C and quenched with 60 µl of ice-cold quench buffer.

To study the effect of nucleosome binding to ISWI, 2.5 µl of purified ISWI, 2.5 µl of nucleosome and 1 µl of protonated buffer were incubated for 30 min on ice and then labeling was performed by adding 54 µl (10 fold) of deuterated labelling buffer for 10 s to 60 minutes, giving final ISWI concentration of 830 nM. The ISWI to nucleosome ratio was kept to 1:1 (830 nM ISWI : 830 nM nucleosome). Previously it has been reported that ISWI binds to the nucleosome substrate with a Kd= 1.3 ± 0.6 nM (64) and as a monomer (21,22,66)). Biochemical as well as structural studies have also shown that ATP/ATP-analogue is not required for binding of ISWI to nucleosome (21, 22, 65, 66)). The labeling reactions was performed at 26°C followed by quenching using ice cold quench buffer (60 µl). This experiment was performed twice.

To study the conformational landscape of ISWI during nucleosome sliding, sliding assays were performed in $D_2O$ sliding/labelling buffer (HEPES pH7.6, 50 mM NaCl, 0.1 mM $MgCl_2$, 0.1 mM EDTA, 1 mM DTT). 2.5 µl of ISWI, 2.5 µl of nucleosome and 1µl of ATP were added to 54 µl of $D_2O$ sliding/labelling buffer. These buffer conditions have been used previously to carry nucleosome remodeling assays (11,22, *66*, *67)* . Concentration of ISWI, nucleosome and ATP were 830 nM, 830 nM and 1.5 mM, respectively. Sliding/labelling was performed for 10s to 60 minutes. The sliding reactions were performed at 26°C and quenched with ice cold quench buffer (60 µl).

**Mass spectrometry**

All quenched samples were injected into a nanoACQUITY UPLC HD/X manager and digested in-line using an immobilized pepsin cartridge (Poros pepsin 2.1 mm D x 30 mm L, Life technologies) at a flowrate 40 µl/min at 15 °C. The peptides were trapped and desalted on an ACQUITY UPLC C18 BEH VanGuard precolumn (1.7 um, 2.1 mm x5 mm, Waters Corp., USA) for 5 min at a flow rate of 40 ul/ min and separated in 10 min using a 10%–90% gradient of acetonitrile in 0.1% formic acid pH 2.5 on a ACQUITY UPLC BEH C18 analytical column (1.7 µm, 1 X 100 mm, Waters Corp., USA) at a flow rate of 40 µl/min at 4°C. Both pre- and analytical columns were in the nanoACQUITY UPLC HD/X manager maintained at 4°C. The eluted peptic peptides were directly analyzed on a Synapt G2 mass spectrometer equipped with a standard ESI source (Waters Corp., USA) over a mass range of 100-2000 m/z. The calibration of the instrument was maintained <3 ppm by continuously spraying Leucine enkephalin (Waters) through the lock mass channel. The mass spectrometer was run in MSE continuous mode with the following settings: ESI positive ion mode; capillary voltage, 3000 V; cone voltage, 40 V; desolvation temperature, 150°C; source temperature, 80°C; nitrogen desolvation gas flow 600 l $hr^{-1}$, scan rate of 0.5 scans $s^{-1}$.

**HDX-MS data analysis**

Peptic fragment identification was performed using Protein Lynx Global Server 2.4 (Waters Corp., USA) from 4-5 MSE analyzed replicates. MSE was performed by a series of low-high collision energies ramping from 18–40 V, ensuring proper fragmentation of the peptic peptides. A sequence coverage of 57% and 52% was obtained for full length ISWI and ATPase domain, respectively (Fig. S18). Alpha fold structure predicts that about 13% of sequence of ISWI is unstructured and usually coverage is not obtained for unstructured regions possibly due to over digestion. Excluding unstructured regions there is about 70% of sequence coverage of FL-ISWI. Only peptides detected across all experiments (ISWI alone as well in presence of ATP and or nucleosome) were analyzed (Fig. S18), however, under any given condition for example in ISWI alone more coverage (73%) and higher peptide degeneracy is observed. Furthermore, we did not observe any increase in coverage by including 3M urea in the quench buffer. Deuterium exchange levels were determined by DynamX 3.0 software (Waters Corp., USA) by identifying the isotopic distribution (from +1 to +6 charge state, depending on the peptide). The parameters used in DynamX 3.0 were: minimum intensity 1000, minimum sequence length 3, maximum sequence length 25, minimum products per amino acid 0.2, Maximum MH+ error 8 ppm and file threshold 3. Isotope distribution and peak selection was verified manually for all peptides. The relative deuterium incorporation was calculated by subtracting the centroid of the isotopic distribution for peptide ions of the unlabeled reference from the centroid of the isotopic distribution for peptide ions from each deuterated sample. All the experiments were performed under identical experimental conditions and hence, deuterium levels are uncorrected for back exchange and are reported as relative fractional uptake (RFU), which is the fraction of total number of exchangeable hydrogens in a peptide (38, 64). Each experiment was performed three times unless stated otherwise.

We obtain only few same peptic peptides for ATPase domain in from ISWI-FL and deletion variant ATPase domain. To compare the deuterium uptake of ATPase domain with corresponding region in ISWI-FL, average deuterium uptake in 50 residue consecutive bins was calculated and plotted along the protein sequence. Comparison at longer bin size did not affect the results.

Correlation plots between deuterium up take of ISWI in presence and absence of sliding were plotted in Origin-9, fitted to a straight-line equation and $r^2$ was taken as conformational correlation value. These values were plotted as heat map. Similarly, correlation plots between deuterium up take of apo-ISWI at different labeling times were compared with each other and r values were plotted as heat map in figure 1. Clustering was performed using R.

**Statistics and reproducibility:** Each experiment was carried three times unless otherwise mentioned, with proteins purified from different batches. Error bars show standard deviation from three independent experiments or spread from two independent experiments as mentioned.

**Competing financial interests:** We declare no competing financial interests.

**Data and material availability:** Any material or data generated in this study will be available upon request to corresponding author.

**Funding:**

This work was supported by an Early Career Research (ECR/2016/000038) award from SERB, DST-GoI to AHW. AHW is recipient of Wellcome Trust-DBT Intermediate Fellowship. The Department of Biotechnology, UoK is funded by DBT-GoI and DST-FIST from DST-GoI. YAB has a senior research fellowship from CSIR-GoI. JYB is recipient of Ramalingaswami Fellowship from DBT-GoI. CIRI is supported by PURSE-grant from DST-GoI. JBU is the recipient of a JC Bose National Fellowship from the GoI.

**Author contributions:**

YAB, JYB & AHW data curation; YAB, JYB, JBU & AHW formal analysis YAB, JYB, JBU & AHW validation; YAB, JYB & AHW investigation; YAB, JYB & AHW visualization; YAB, JYB & AHW methodology; AHW, YAB, JBU & JYB writing-original draft; AHW, JBU & YAB conceptualization; AHW (Early Career Research (ECR/2016/000038) award from SERB) funding acquisition; AHW, YAB, JYB, JBU & SA writing-review and editing; AHW & JBU supervision; AHW project administration

| | Start-residue | End-residue | RFU at 10s of labeling | RFU at 1 hour of labeling | Difference (1 hour - 10s) |
|---|---|---|---|---|---|
| **Class-a** | 427 | 436 | 0.0291 | 0.0326 | 0.0035 |
| | 425 | 436 | 0.024 | 0.048 | 0.024 |
| | 365 | 371 | 0.0352 | 0.0634 | 0.0282 |
| | 365 | 377 | 0.0193 | 0.0646 | 0.0453 |
| **Class-b** | 672 | 680 | 0.4042 | 0.3602 | -0.044 |
| | 684 | 696 | 0.3199 | 0.3065 | -0.0134 |
| | 686 | 696 | 0.3392 | 0.3318 | -0.0074 |
| | 586 | 593 | 0.3899 | 0.3858 | -0.0041 |
| | 253 | 270 | 0.2897 | 0.292 | 0.0023 |
| | 641 | 648 | 0.317 | 0.3206 | 0.0036 |
| | 627 | 640 | 0.4856 | 0.4936 | 0.008 |
| | 672 | 683 | 0.3892 | 0.3984 | 0.0092 |
| | 104 | 112 | 0.3954 | 0.4095 | 0.0141 |
| | 608 | 621 | 0.2737 | 0.2888 | 0.0151 |
| | 665 | 671 | 0.3148 | 0.3304 | 0.0156 |
| | 282 | 295 | 0.2886 | 0.3046 | 0.016 |
| | 11 | 35 | 0.3158 | 0.3321 | 0.0163 |
| | 253 | 268 | 0.2629 | 0.2829 | 0.02 |
| | 649 | 664 | 0.4139 | 0.4359 | 0.022 |
| | 256 | 268 | 0.3191 | 0.3438 | 0.0247 |
| | 608 | 620 | 0.2904 | 0.3156 | 0.0252 |
| | 209 | 218 | 0.2646 | 0.2953 | 0.0307 |
| | 641 | 650 | 0.3874 | 0.4252 | 0.0378 |
| **Class-c** | 505 | 516 | 0.3996 | 0.4555 | 0.0559 |
| | 167 | 191 | 0.0983 | 0.1593 | 0.061 |
| | 660 | 671 | 0.082 | 0.1566 | 0.0746 |
| | 36 | 50 | 0.2104 | 0.2851 | 0.0747 |
| | 707 | 727 | 0.4687 | 0.5435 | 0.0748 |
| | 37 | 50 | 0.216 | 0.2913 | 0.0753 |
| | 626 | 640 | 0.4548 | 0.5312 | 0.0764 |
| | 576 | 593 | 0.3516 | 0.4302 | 0.0786 |
| | 383 | 394 | 0.4951 | 0.5759 | 0.0808 |
| | 624 | 640 | 0.3977 | 0.4799 | 0.0822 |
| | 123 | 130 | 0.2731 | 0.3646 | 0.0915 |
| | 124 | 130 | 0.2943 | 0.3863 | 0.092 |
| | 707 | 728 | 0.4059 | 0.5007 | 0.0948 |
| | 152 | 163 | 0.321 | 0.4166 | 0.0956 |
| | 574 | 585 | 0.3577 | 0.4661 | 0.1084 |
| | 944 | 950 | 0.2963 | 0.4054 | 0.1091 |

| | | | | | |
|---|---|---|---|---|---|
| | 381 | 394 | 0.4605 | 0.5706 | 0.1101 |
| | 271 | 281 | 0.1541 | 0.265 | 0.1109 |
| | 553 | 560 | 0.0324 | 0.1449 | 0.1125 |
| | 380 | 394 | 0.4387 | 0.5519 | 0.1132 |
| | 494 | 501 | 0.202 | 0.3197 | 0.1177 |
| | 378 | 394 | 0.3751 | 0.4962 | 0.1211 |
| | 536 | 552 | 0.1401 | 0.2627 | 0.1226 |
| | 437 | 453 | 0.0361 | 0.1606 | 0.1245 |
| | 411 | 426 | 0.1858 | 0.3169 | 0.1311 |
| | 117 | 130 | 0.2746 | 0.4076 | 0.133 |
| | 943 | 950 | 0.2174 | 0.3529 | 0.1355 |
| | 51 | 60 | 0.1575 | 0.3023 | 0.1448 |
| | 115 | 130 | 0.2289 | 0.3749 | 0.146 |
| | 561 | 569 | 0.0302 | 0.1833 | 0.1531 |
| | 437 | 450 | 0.0479 | 0.2031 | 0.1552 |
| | 746 | 760 | 0.2113 | 0.3757 | 0.1644 |
| | 600 | 607 | 0.1704 | 0.3398 | 0.1694 |
| | 1011 | 1025 | 0.1635 | 0.3429 | 0.1794 |
| | 300 | 307 | 0.0334 | 0.215 | 0.1816 |
| | 239 | 252 | 0.0793 | 0.2666 | 0.1873 |
| | 478 | 493 | 0.087 | 0.2826 | 0.1956 |
| | 329 | 340 | 0.0093 | 0.2066 | 0.1973 |
| | 478 | 492 | 0.1229 | 0.3222 | 0.1993 |
| | 809 | 822 | 0.0646 | 0.2795 | 0.2149 |
| | 793 | 808 | 0.1008 | 0.331 | 0.2302 |
| | 784 | 792 | 0.03 | 0.262 | 0.232 |
| | 786 | 792 | 0.024 | 0.2565 | 0.2325 |
| | 333 | 340 | 0.0249 | 0.2619 | 0.237 |
| | 764 | 783 | 0.2015 | 0.4523 | 0.2508 |
| | 823 | 834 | 0.134 | 0.4202 | 0.2862 |
| | 819 | 834 | 0.1142 | 0.4965 | 0.3823 |
| | 568 | 575 | 0.027 | 0.4952 | 0.4682 |

**Supplementary Table 1:** Showing classification of different regions of ISWI based on their deuterium exchange in the resting-state.

Fig. S1

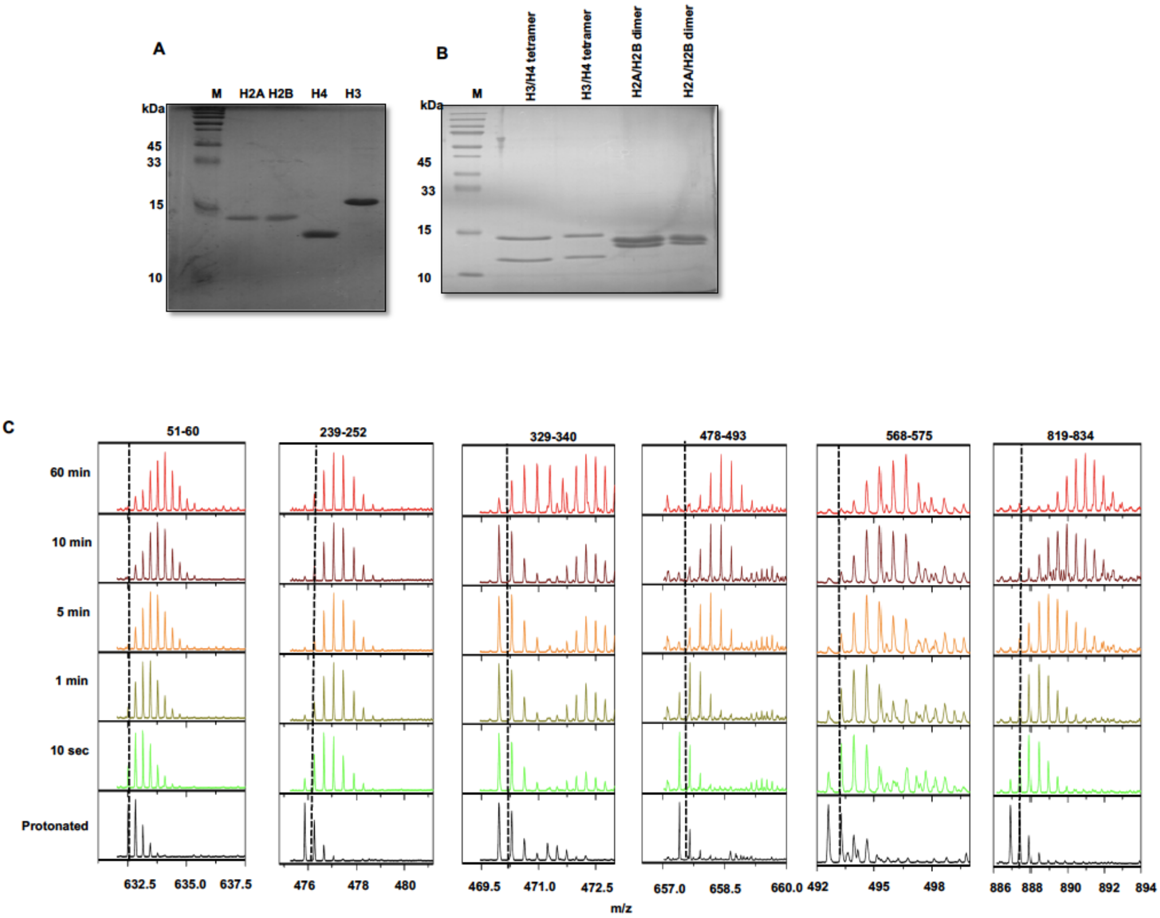

**Supplementary Figure 1: Protein purification:** A) SDS-PAGE of purified histones. (B) SDS-PAGE showing purified histone H2A-H2B dimer and H3-H4 tetramer. (C) Representative mass spectra of different regions of ISWI labeled for different times. Doted lines show the shift in centroid of mass relative to protonated protein.

**Fig. S2**

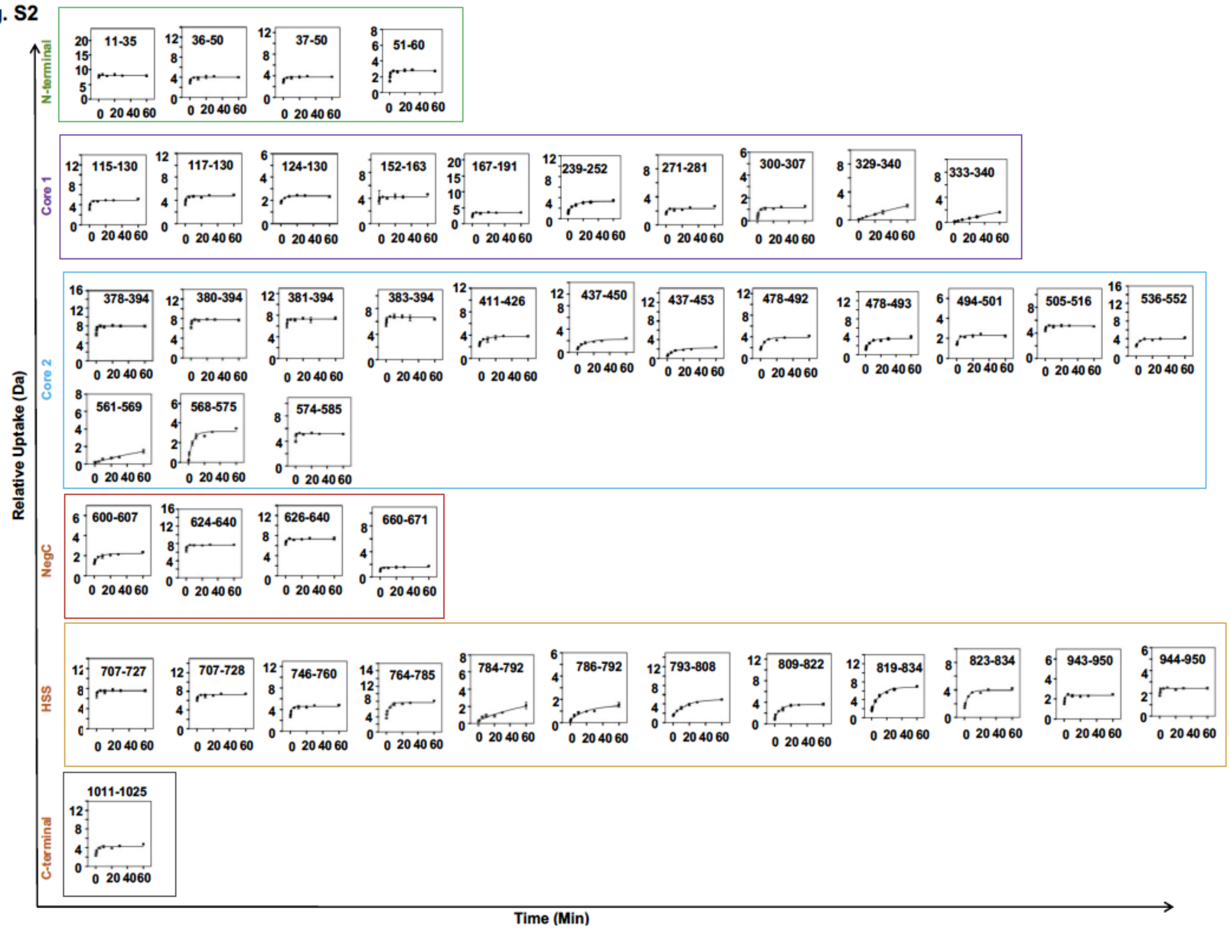

**Supplementary Figure 2: Deuterium uptake due to intrinsic fluctuations of ISWI:** Deuterium uptake by different regions of ISWI relative to protonated protein is plotted against different labelling times. Uptake curves are fitted to single exponential function. Numbers inside the graphs denote the starting and end residue of peptide. Error bars show standard deviation from three different experiments.

Fig. S3

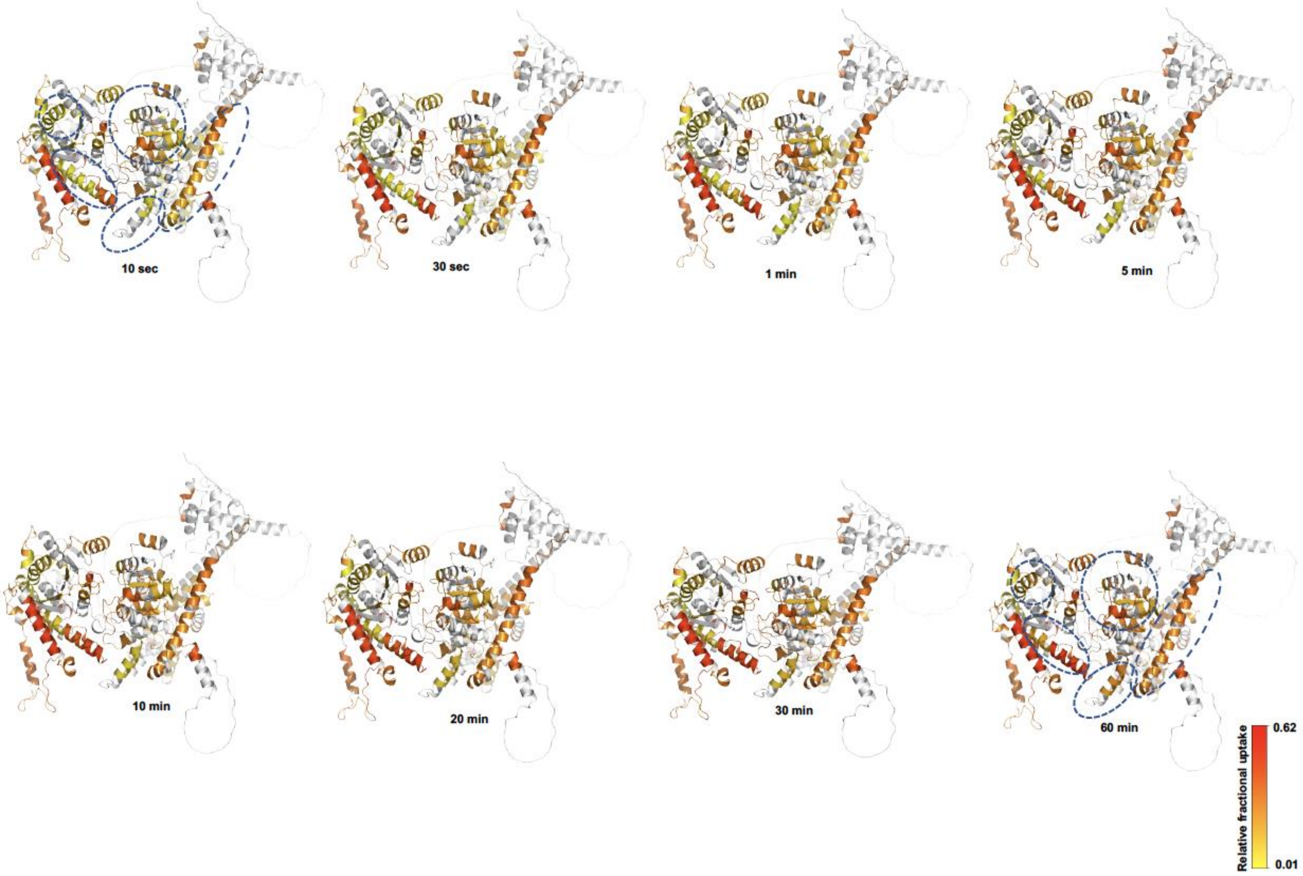

**Supplementary Figure 3: Conformations sampled intrinsically by ISWI:** RFU values of different regions at different labelling times were mapped on to 3D structure of ISWI.

Fig. S4

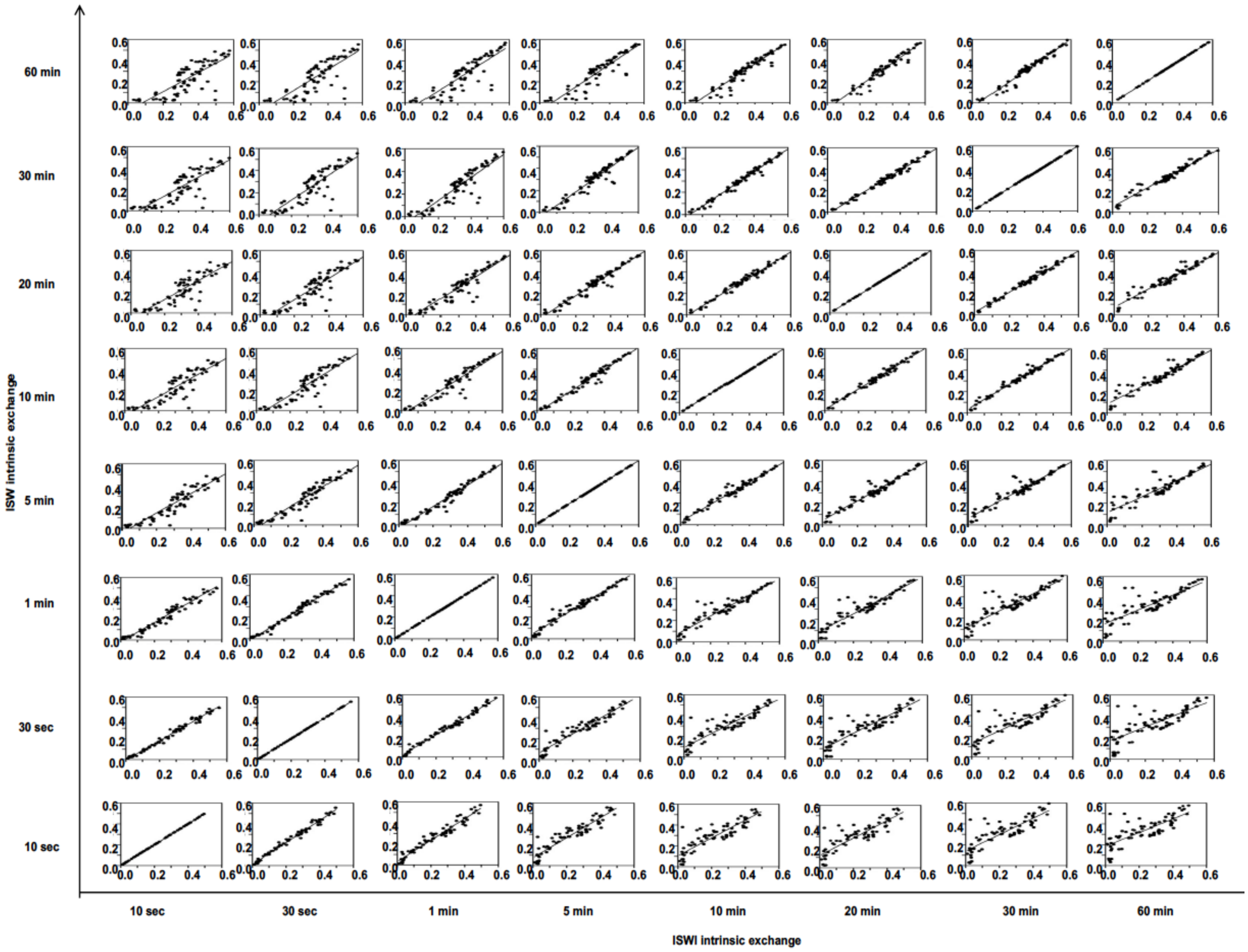

**Supplementary Figure 4: Conformational correlation:** Correlation plots between deuterium up take of corresponding regions of ISWI obtained at different labeling times.

**Fig. S5**

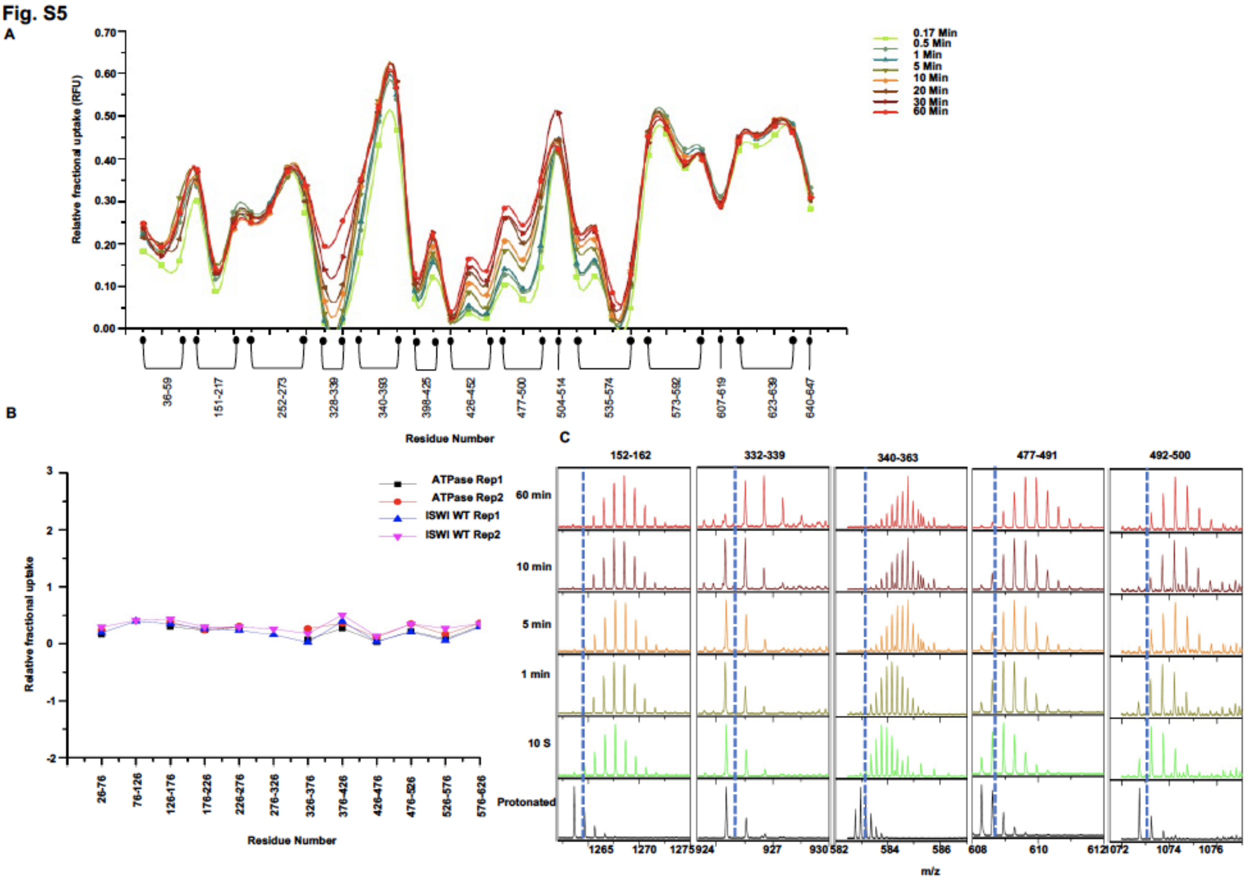

**Supplementary Figure 5: Deuterium uptake by ATPase domain:** (A) RFU of ATPase domain exchanged for different times (10 s - 60 min) (B) RFU values of ATPase domain and corresponding region of ISWI-FL. We did not obtain exactly same peptide for ATPase domain as that of the corresponding region in ISWI-FL. Deuterium uptake values were binned for every consecutive 50 residue bins to compare deuterium uptake profile between ATPase domain and ISWI-FL. (C) Representative mass spectra of peptides from ATPase domain exchanged for different time points. Doted lines show the centroid of mass spectra for protonated protein.

Fig. S6

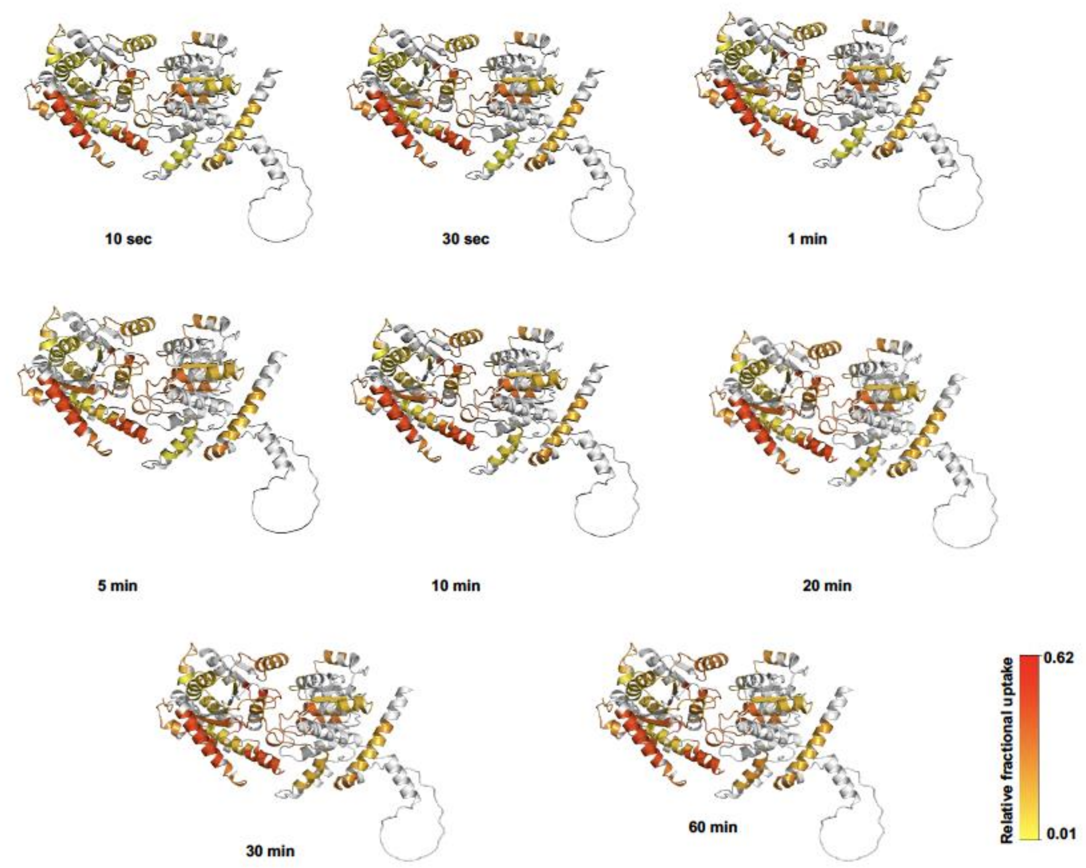

**Supplementary Figure 6: Conformations sampled by ATPase domain:** RFU values of different regions at different labelling times were mapped on to 3D structure of ATPase domain.

**Fig. S7**

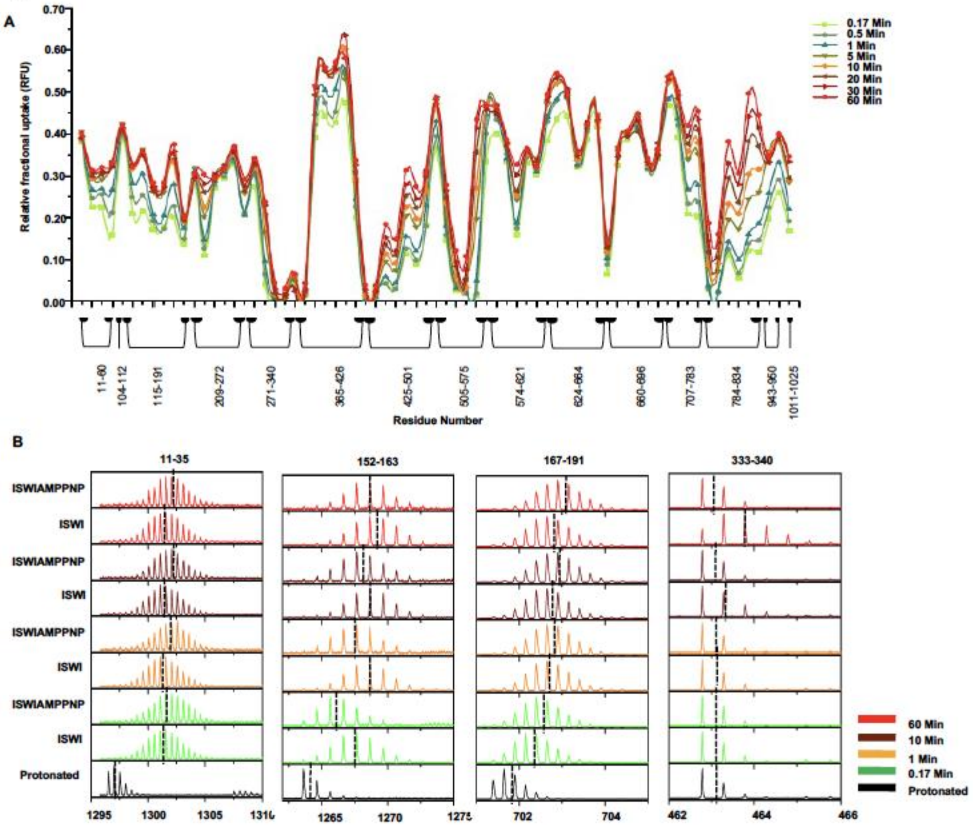

**Supplementary Figure 7: Deuterium uptake by ISWI in presence of AMPPNP:** (A) RFU values of ISWI-FL in presence of ATP analog, AMPPNP. (B) Representative mass spectra of different peptides of ISWI in presence and absence of AMPPNP and exchanged for different times. Doted lines show the centroid of mass spectra.

Fig. 8

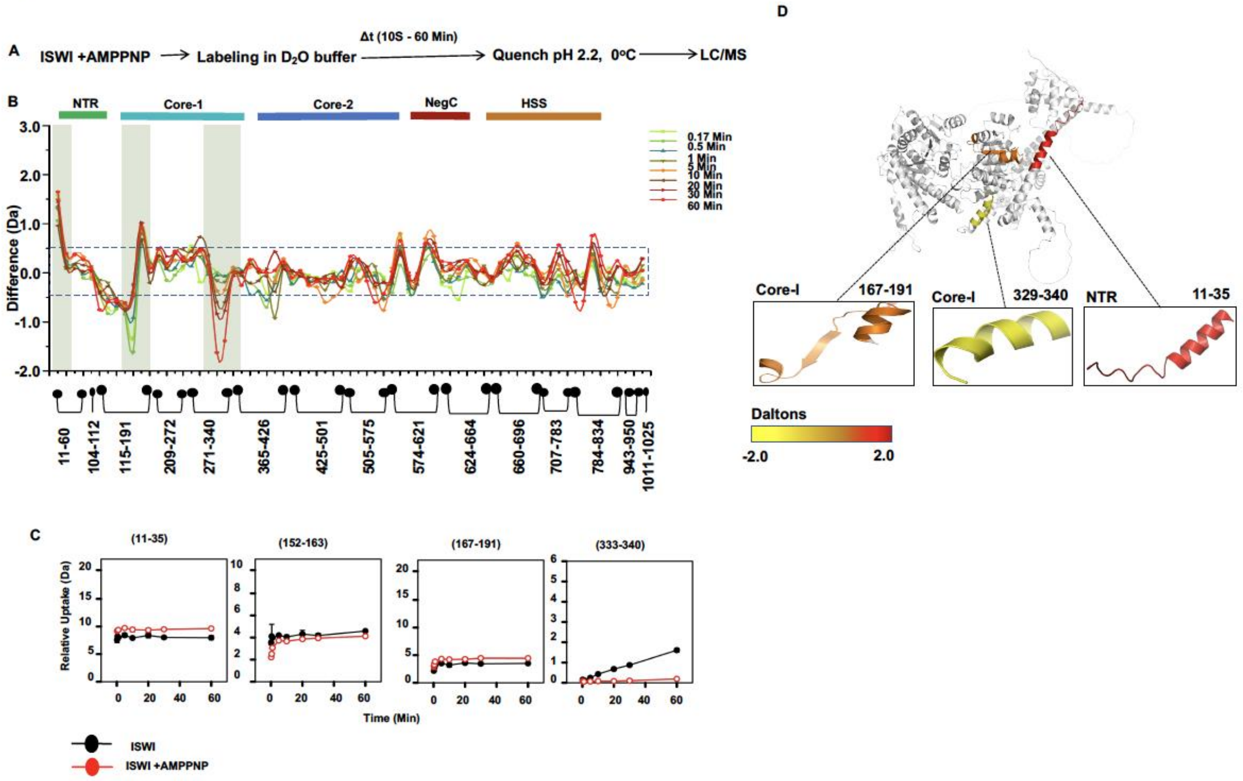

**Supplementary Figure 8. Effect of AMPPNP binding on intrinsic dynamics of ISWI:** (A)Flow chart showing the main steps of the HDX-MS experiment for ISWI-AMPPNP interaction. (B) Difference (subtraction) plot of ISWIAMPPNP and ISWI at different time points of labelling. Differences in deuterium exchange of more than $\pm 0.5$ Da are considered significant (dashed box). Shaded regions show significant change. (C) Representative uptake plots of different regions of ISWI in presence and absence of AMPPNP. (D) Regions of ISWI showing significant differential deuterium uptake in the presence of AMPPNP at 60 min of labeling are mapped on to the Alpha fold structure of ISWI.

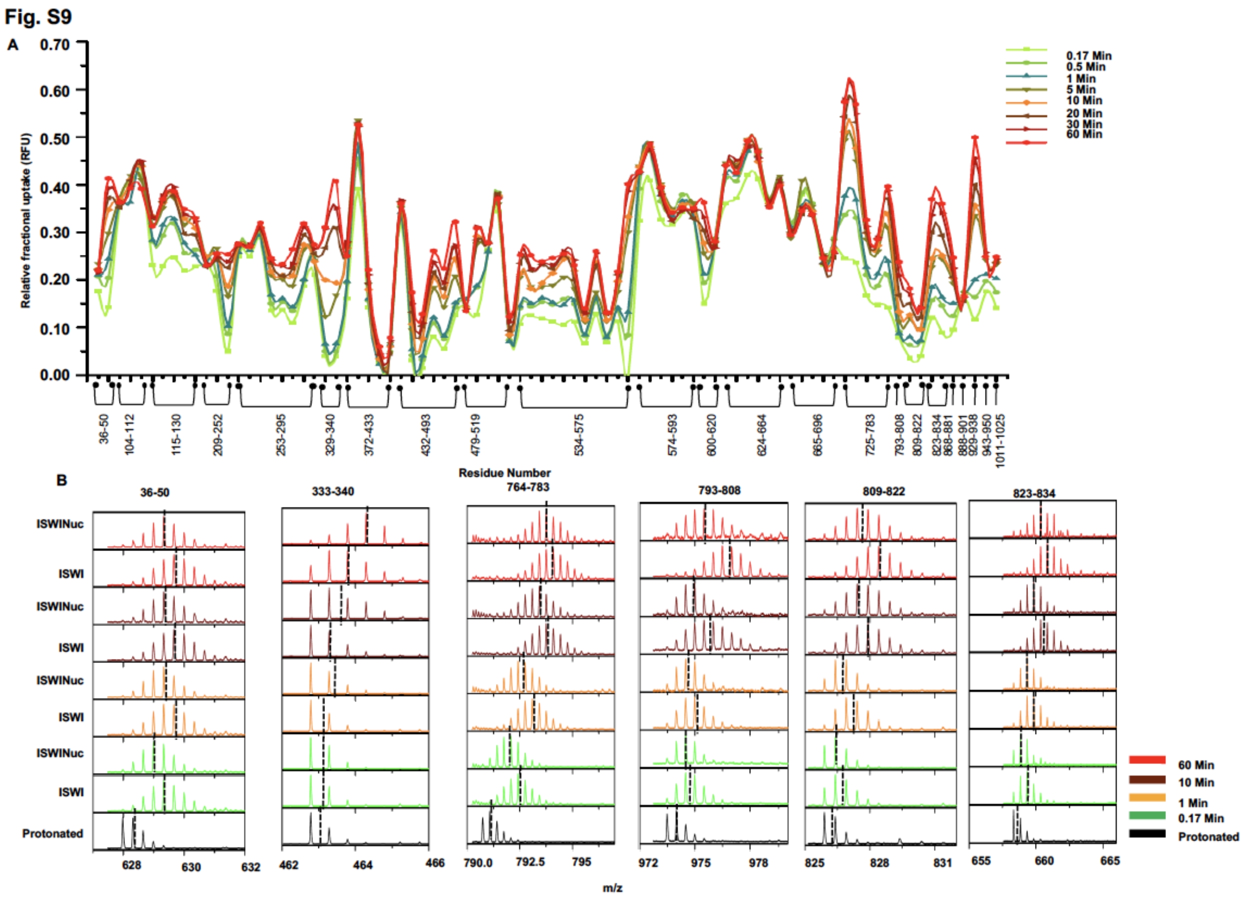

**Supplementary Figure 9: Deuterium uptake by ISWI in presence of nucleosome:** (A) RFU values for ISWI-FL in presence of nucleosome obtained after different times of labelling (10 s - 60 min). Each dot represents one peptic fragment. (B) Representative mass spectra of different peptides of ISWI protein in presence and absence of nucleosome exchanged for different times. Doted lines show the centroid of mass spectra.

**Fig. S10**

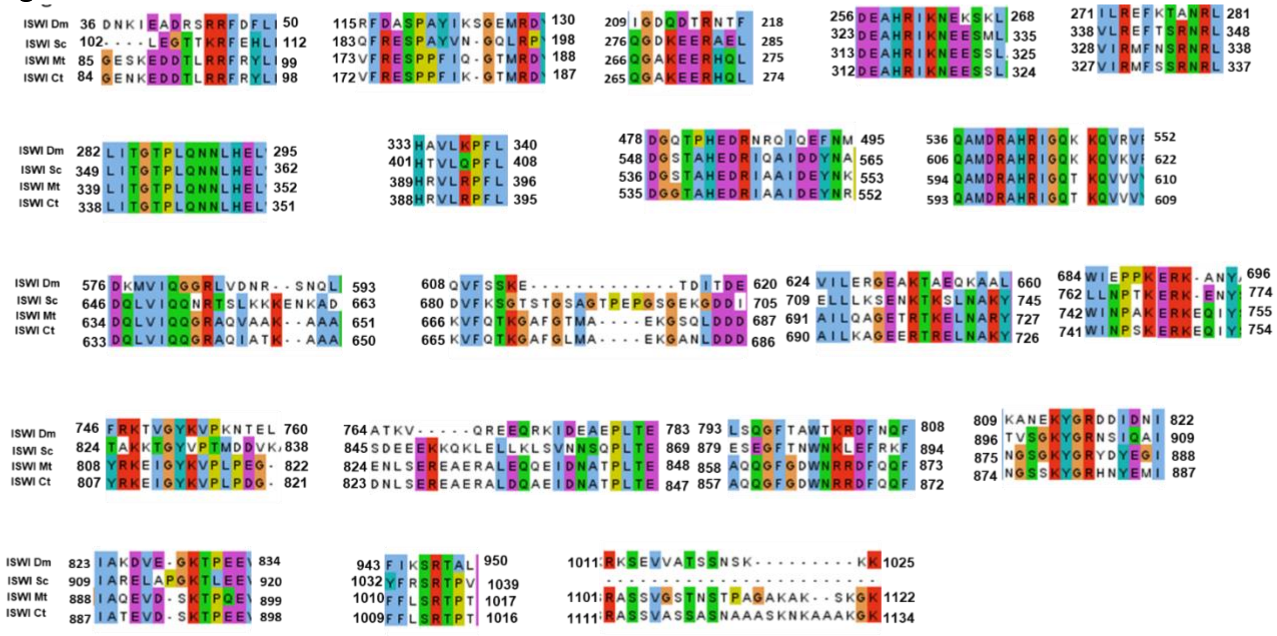

**Supplementary Figure 10: Sequence conservation of regions of ISWI undergoing conformational change during nucleosome sliding:** Sequence conservation of regions of ISWI involved in nucleosome sliding determined by Clustal X sequence alignment. blue-hydrophobic, red- positive charge, magenta-negative charge, green-polar, pink-cysteines, orange-glycine, yellow-prolines, cyan-aromatic and white unconserved amino acids.

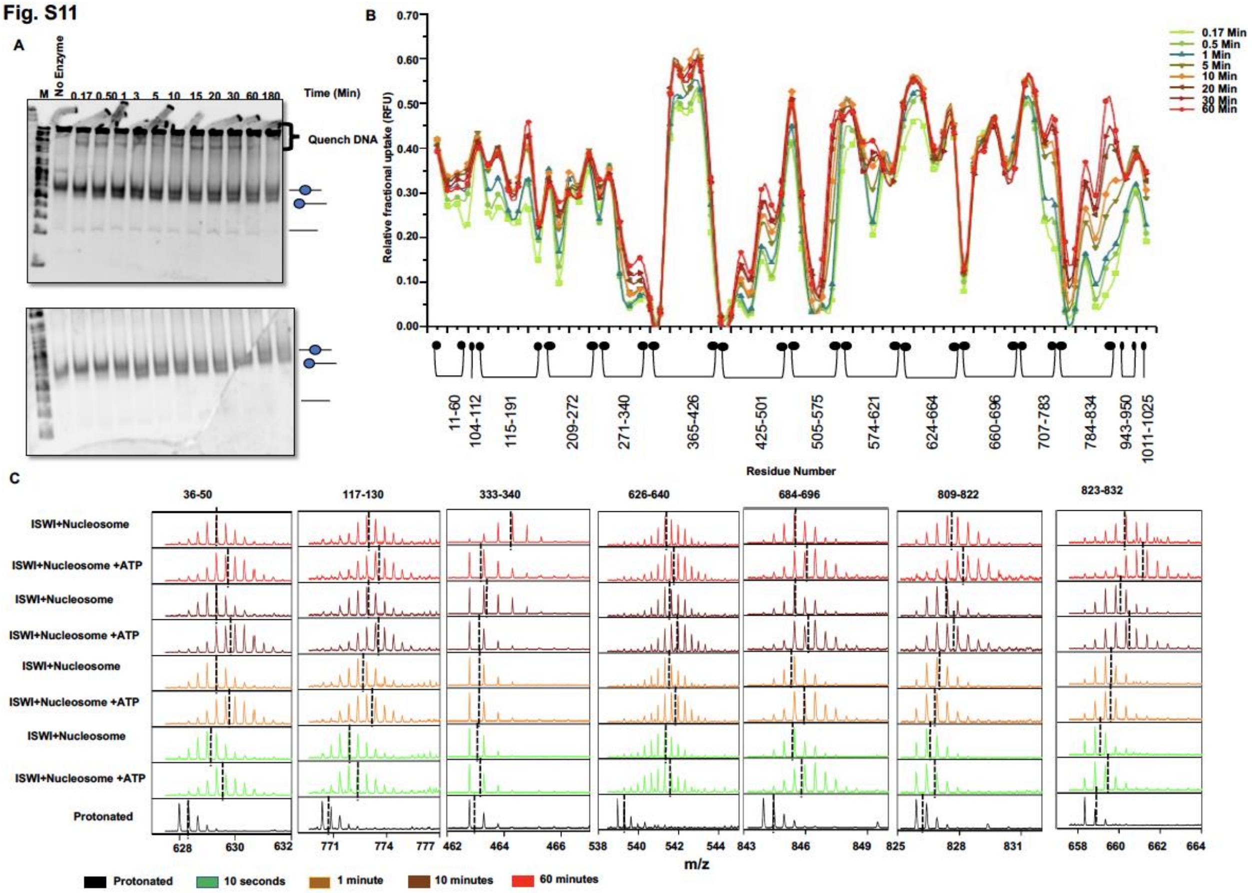

**Supplementary Figure 11: Deuterium uptake by ISWI during nucleosome sliding:** (A) Native-PAGE gel images corresponding to image in figure 4A. (B) RFU values for ISWI during mono-nucleosome sliding performed in deuterated buffer for different time durations (10 s - 60 min). (C) Representative mass spectra of different peptides ISWI in nucleosome-bound state or during nucleosome sliding. Doted lines show the centroid of mass spectra.

Fig. S12

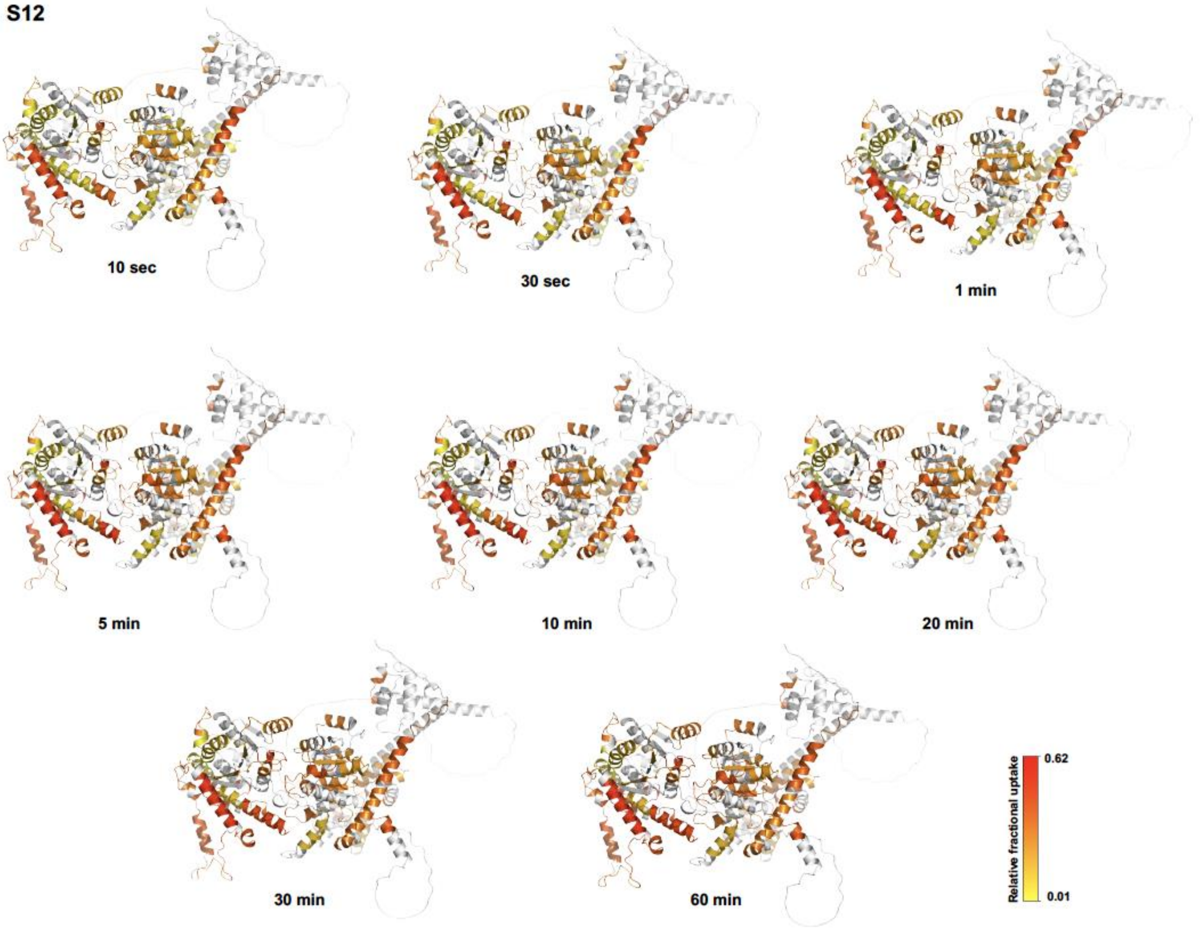

**Supplementary Figure 12: Conformations adopted by ISWI during nucleosome sliding:** RFU values of different regions at different sliding/labelling times were mapped on to 3D structure of ISWI.

Fig. S13

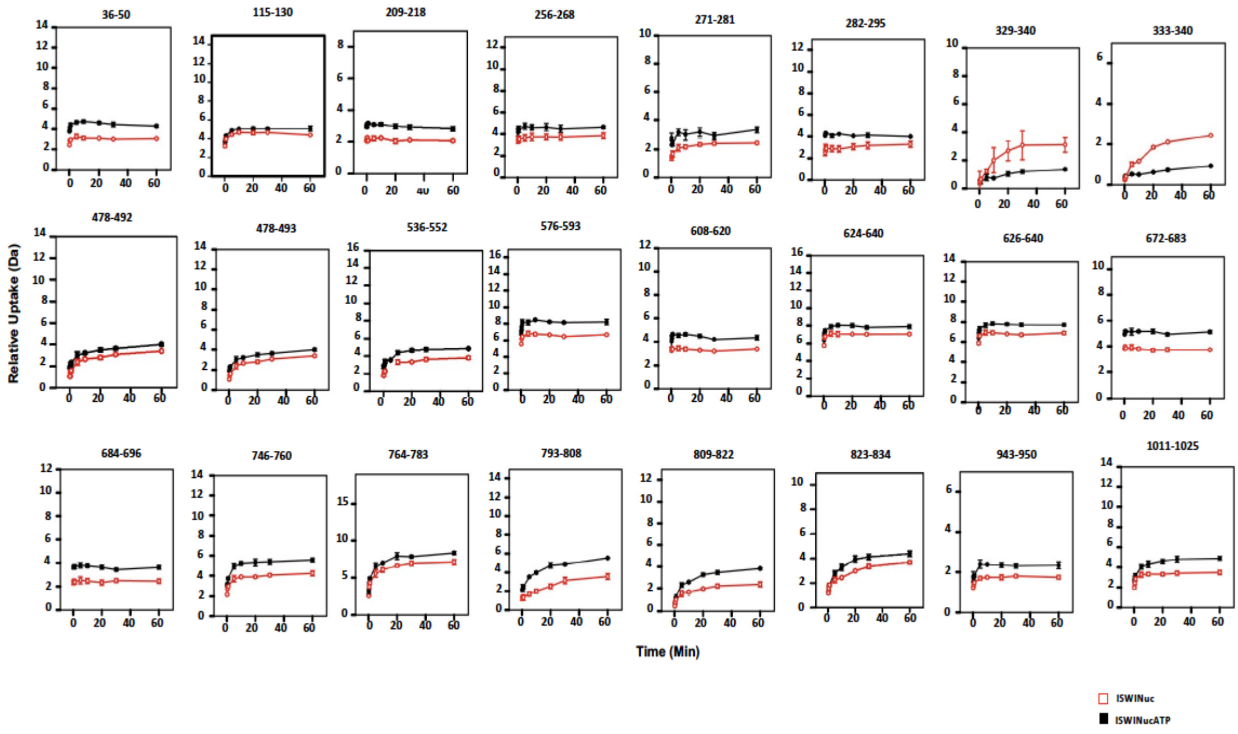

**Supplementary Figure 13: Deuterium uptake by ISWI during nucleosome sliding and in nucleosome-bound state:** Relative Deuterium uptake plots of different regions of ISWI during sliding (black) and nucleosome binding (red).

Fig. S14

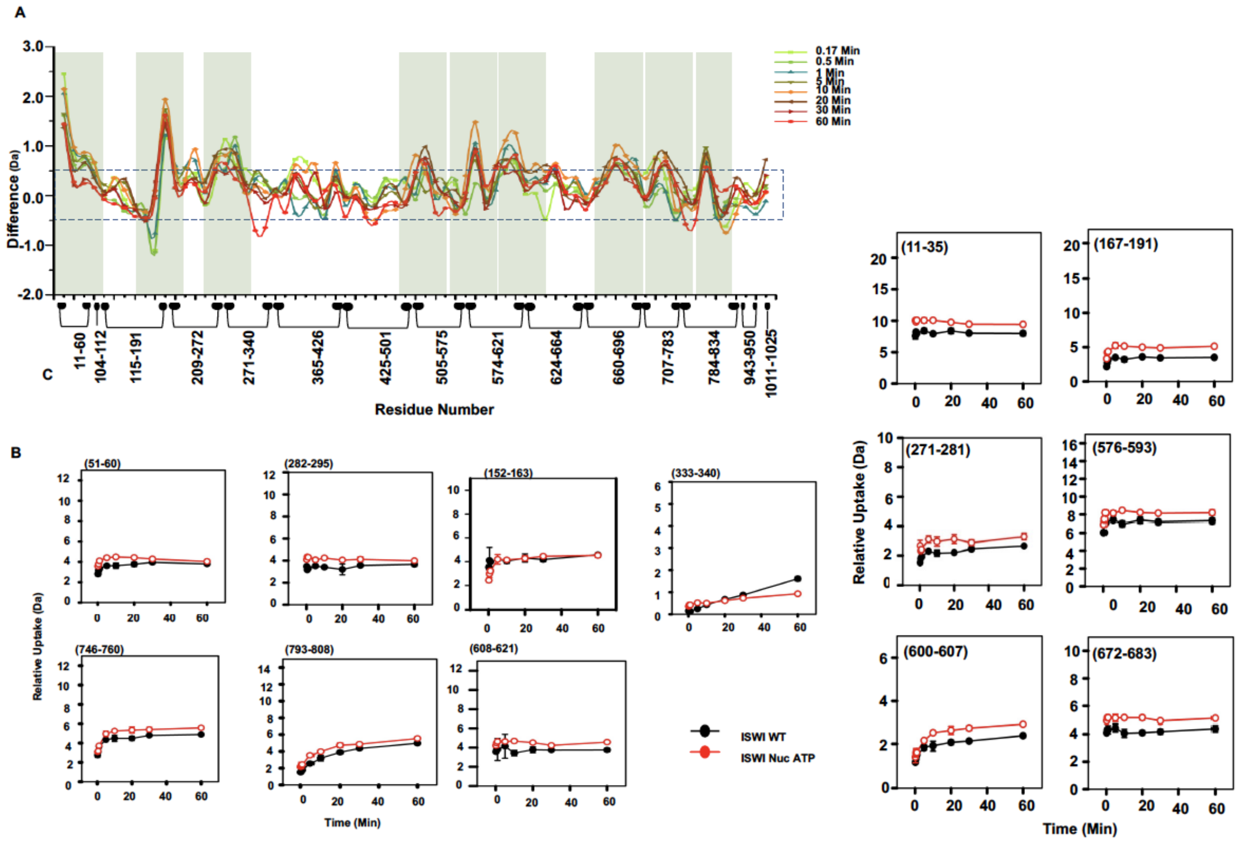

**Supplementary Figure 14. Conformational dynamics of ISWI during nucleosome sliding:** (A)Difference (subtraction) plot of ISWI during nucleosome sliding and ISWI alone at different labelling times. Differences in deuterium exchange more than ±0.5 Da are considered significant (dashed box). Shaded regions show regions with significant change. (B) Representative deuterium uptake plots of different regions of ISWI during nucleosome sliding (red) or alone (black) obtained from three different experiments. Error bars show the standard deviations.

Fig. S15

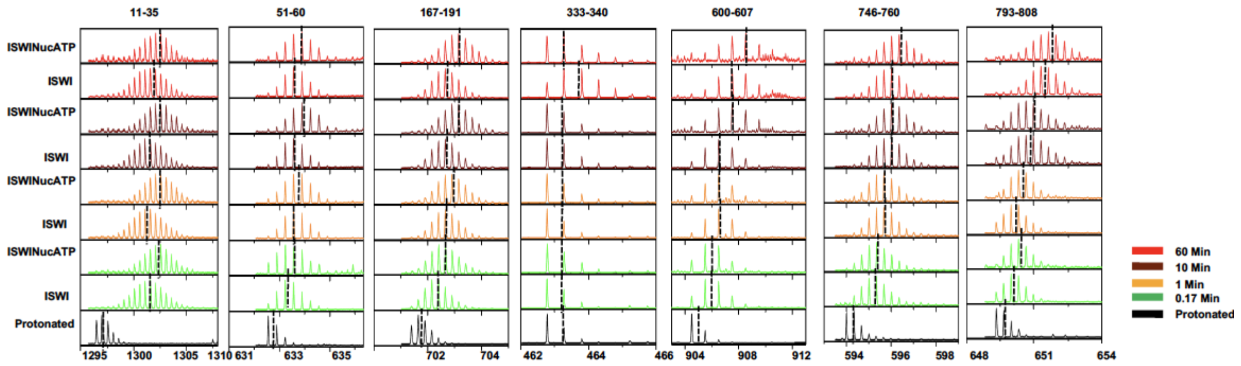

**Supplementary Figure 15: Deuterium uptake by ISWI during nucleosome sliding:** Representative mass spectra of different peptides of ISWI in apo-state or during nucleosome sliding. Doted lines show the centroid of mass spectra.

**Fig. S16**

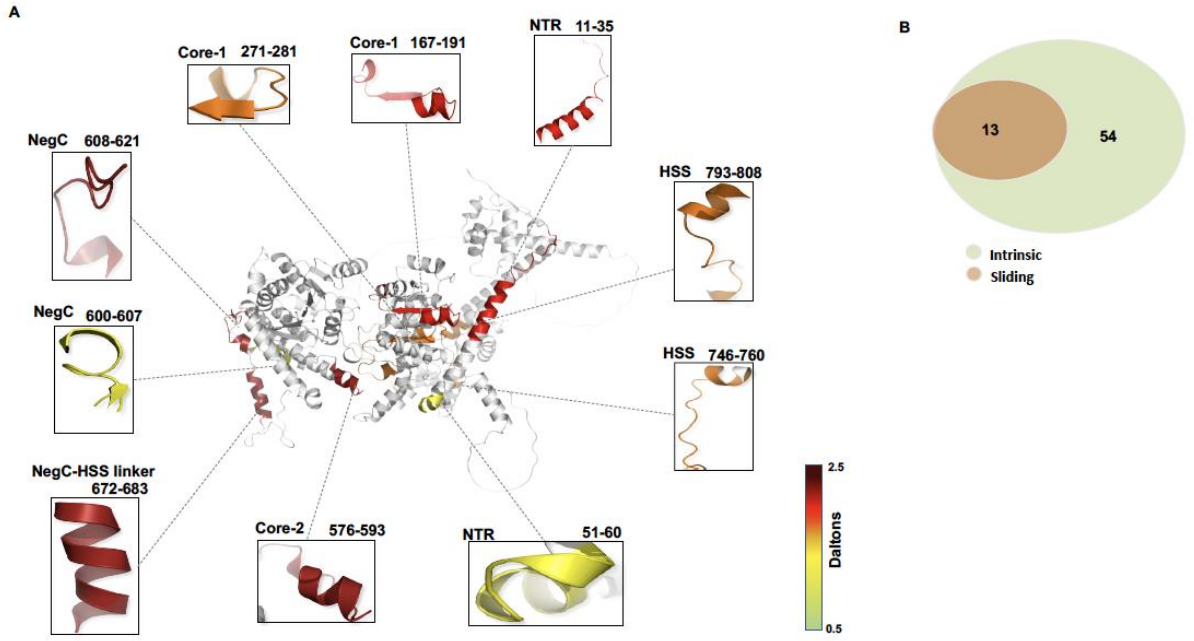

**Supplementary Figure 16. Regions of ISWI undergoing conformational change during nucleosome sliding:** (A) Regions of ISWI showing significant differential deuterium uptake during nucleosome sliding relative to intrinsic uptake at 10 min of labeling are mapped on to the ISWI Alpha fold structure. (B) Overlap between regions showing increased deuterium uptake during nucleosome sliding (in comparison to apo-ISWI) and intrinsically fluctuating regions from Class-b and Class-c.

**Fig. S17**

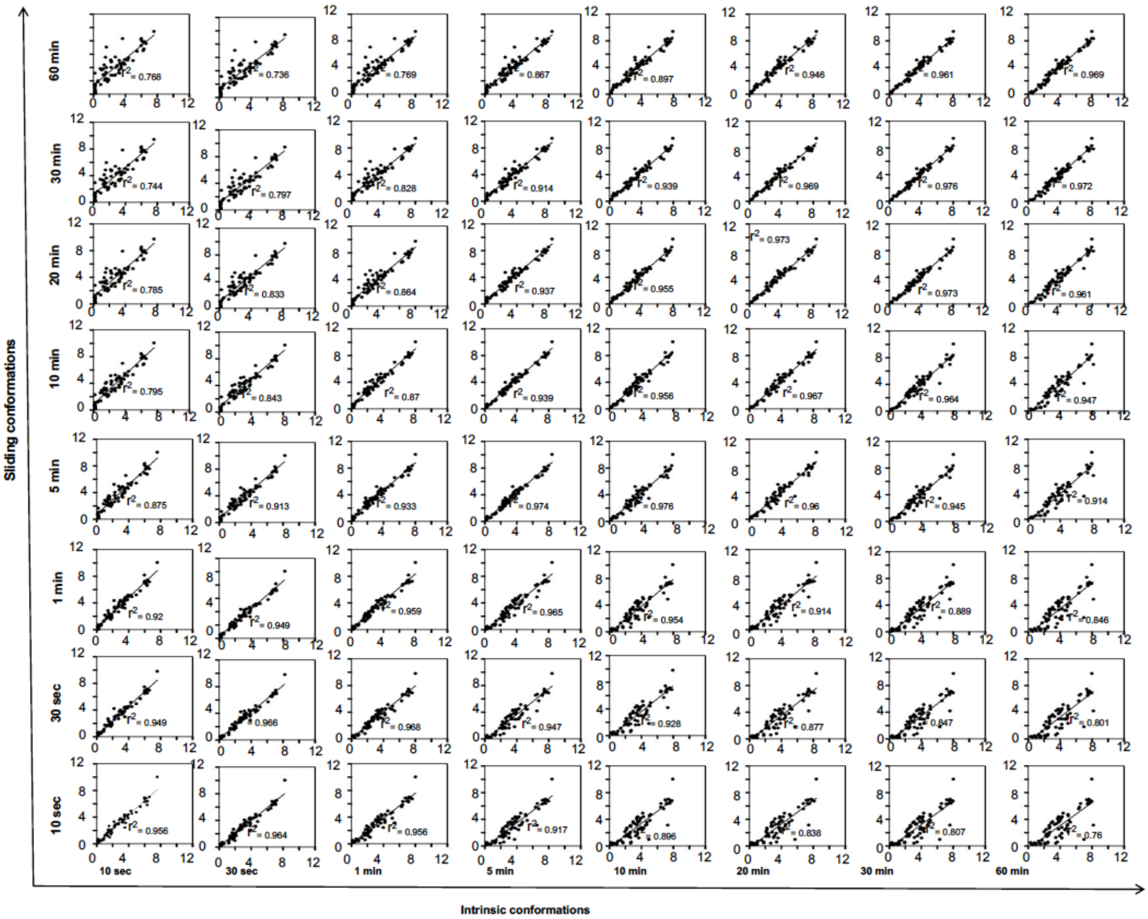

**Supplementary Figure 17: Conformational correlation:** Correlation plots between deuterium up take of corresponding regions of ISWI alone and during nucleosome sliding at same labelling times.

**Fig. S18**

**Coverage map of ISWI**

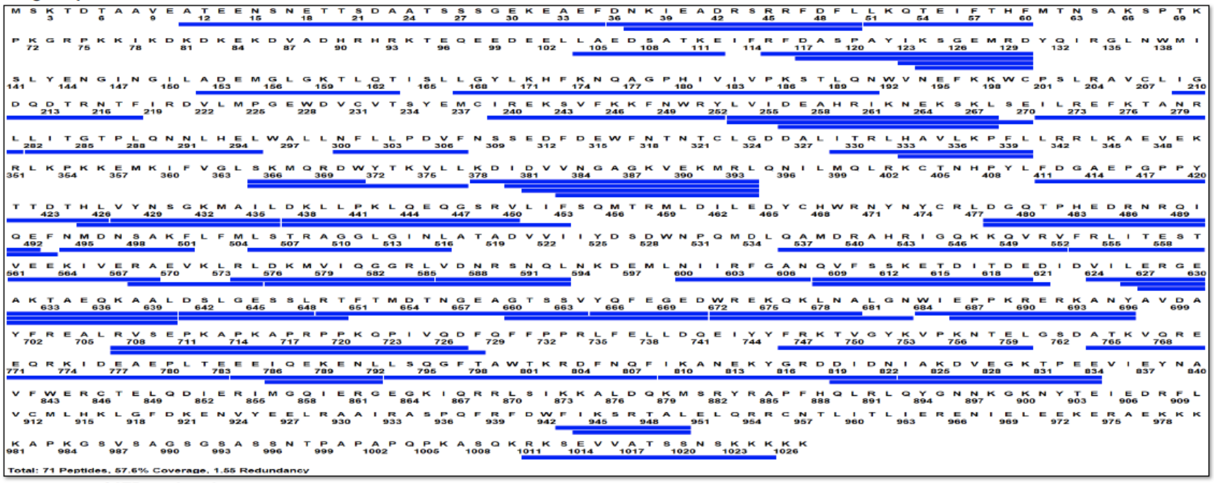

**Coverage map of ATPase domain**

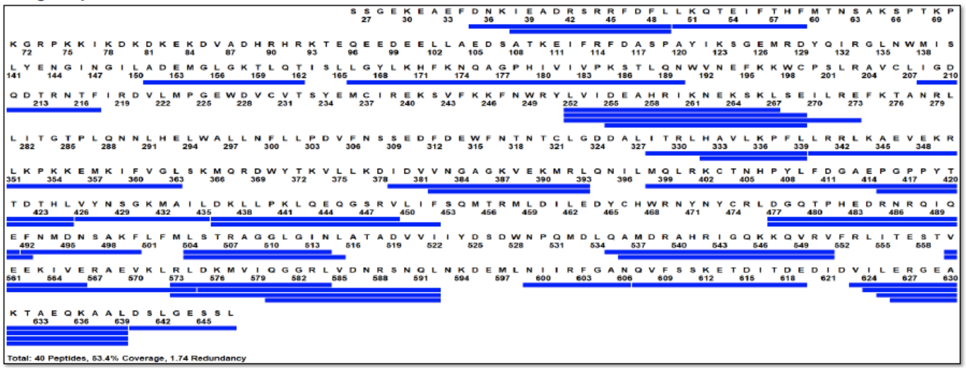

**Supplementary Figure 18: Sequence coverage:** (top) Map showing peptides (blue bars) detected across sequence of FL-ISWI accounting for 57% of coverage or about 70% of coverage excluding unstructured regions predicted by alpha fold. (below) ATPase domain coverage (52%).
